# Elastic tethers remain functional during anaphase arrest in partially-lysed crane-fly spermatocytes: a possible approach for studying mitotic tethers

**DOI:** 10.1101/2024.05.18.594823

**Authors:** Aisha Aidil, Samir A. Malick, Arthur Forer

## Abstract

Mitotic tethers connect partner telomeres of all segregating anaphase chromosomes in all animal cells that have been tested, as detected by laser-cutting chromosome arms during anaphase and seeing that the arm fragments move rapidly across the equator to their partner chromosome moving to the opposite pole, telomere moving towards telomere. Tethers exert anti-poleward forces on the poleward separating telomeres, but tether elasticity (that produces the backwards forces) diminishes during anaphase: as determined by the behavior of arm fragments; short tethers (early anaphase) are elastic, long tethers (late anaphase) are not elastic, and medium-length tethers transition between the two states. We developed a procedure in which the tethers still functioned after we partially-lysed anaphase crane-fly spermatocytes. The partial lysis consistently arrested chromosome movements, after which the tethers moved the chromosomes backwards, potentially allowing the elastic tethers to be studied biochemically. To ensure that tether function was not altered by the partial cell-lysis procedure, we compared backward chromosome movements in partially-lysed cells with arm fragment movements in control cells. In the partially-lysed cells the backward chromosomal movements had characteristics identical to those of arm fragments in non-lysed (control) cells. In particular, in both control and partially-lysed cells shorter tethers caused backward movements more often than did longer tethers; shorter tethers caused backward movements over greater fractional distances (of the tether) than did longer tethers; and velocities of the backwards movements were the same for tethers of different lengths. We also compared the effects of Calyculin A (an inhibitor of Protein Phosphatase1) in control *versus* in partially-lysed cells. Calyculin A (CalA) added to control cells in early anaphase blocks dephosphorylation, thereby maintaining tether elasticity throughout anaphase: after the chromosomes reach the poles they move backwards when the usual poleward forces are reduced. Partial lysis preserves this tether functionality: after partial lysis of CalA-treated cells the chromosomes move backward and reach the partner telomeres at even very long tether lengths. We conclude that partial cell-lysis arrests anaphase chromosome poleward movement but does not affect tether function.

## INTRODUCTION

Our experiments deal with *mitotic tethers*, newly-discovered components of the spindle apparatus. Mitotic tethers are structural connections between the telomeres of all separating anaphase chromosomes in all animal cells so-far tested. Their presence was deduced from experiments using laser microbeam irradiations. The basic experiment was to cut a chromosomal arm in early anaphase; the resultant arm fragment moved backwards in the anti-poleward direction, the arm fragment’s telomere moving at fast speed to its partner telomere in the opposite half spindle, while the amputated chromosome continued moving poleward (LaFountain et al., 2002). The later in anaphase the arm was cut the shorter the backward distance that the arm fragment moved (LaFountain et al., 2002), indicating that tethers become less elastic as anaphase progresses. The original experiments (LaFountain et al., 2002) were done using crane-fly spermatocytes but subsequent experiments using a laser microbeam to cut chromosome arms found the same phenomena in a wide variety of animal cells ranging from aquatic flatworms to insects to spiders to marsupials to humans (Forer *et al.,* 2017). There is some evidence that tethers are involved in coordinating movements between chromosomes (Sheykhani *et al.,* 2017; Forer and Berns, 2020; Fegaras-Arch et al., 2020).

“Tethers”, not spindle microtubules, cause the laser-cut arm fragments of anaphase chromosomes to move backwards. Microtubules do not cause the anti-poleward movement because arm fragment movements require both telomeres to be intact: ablating either of the two partner telomeres stops the arm fragment movement (LaFountain *et al.,* 2002; Forer *et al.,* 2021). Also, after cutting individual arm fragments in half only the sub-fragment with the telomere moves toward the partner telomere while the other sub-fragment stops moving (LaFountain *et al.,* 2002). Further, taxol greatly slows down chromosomal poleward movement compared to control cells while arm fragments move at the same speeds as in control cells (Forer *et al.,* 2018). Therefore, microtubules are not force generators for the backwards movements of arm fragments.

Ultra-fine DNA bridges do not cause arm-fragments to move backwards, either. Ultra-fine bridges sometimes connect partner telomeres, but they are not elastic, are not found in all cells, and when found they connect telomeres of only a small number of the anaphase chromosome pairs; they are mostly at the centromeric regions rather than at telomeres (Chan *et al.,* 2007; Barefield and Karlseder, 2012; Gemble *et al*., 2015), and they retard or stop anaphase segregation (Su *et al*., 2016). This contrasts with mitotic tethers which are present between the telomeres of every separating pair of chromosomes, connect two out of the four arms of each segregating meiotic chromosome pair and one out of the two arms in each segregating mitotic pair. Mitotic tethers do not slow down anaphase segregation since cutting them does not affect anaphase speed (LaFountain *et al*., 2002; Forer *et al.,* 2017; Sheykhani *et al.,* 2017; Forer *et al*., 2018: Forer and Berns, 2020). Thus, neither microtubules nor ultra-fine DNA bridges move the arm fragments.

That tethers are separate structural components connecting telomeres of separating anaphase chromosomes can be seen directly in electron microscope tomograms of crane-fly spermatocytes (Forer and Otsuka, 2023), tethers appearing as two component structures extending between separating partner telomeres.

In sum, mitotic tethers are distinct structural entities that in anaphase extend between separating telomeres. They are neither microtubules nor ultra-fine DNA bridges.

Tethers are elastic in early anaphase and non-elastic by late anaphase as judged by their ability to move arm fragments backward, away from the pole to which they originally were moving. The tethers connecting chromosome arms produce forces on the attached chromosome enough to stretch the arms by about 10% (Forer et al., 2017) but the backward forces from tethers are much smaller than the forward, poleward forces on kinetochores: cutting tethers during anaphase (or cutting all arms) does not change the speeds of kinetochore (chromosome) poleward movement (Sheykhani et al., 2017; Forer and Berns, 2020). Tethers remain attached to the telomeres even in late anaphase, when tethers are not elastic; we know this because directly cutting late anaphase tethers, or cutting arms in late anaphase to produce arm fragments (that do not move), causes the arms (or the non-cut arm) to shrink by about 10%. The overall picture is of tethers being elastic until mid-anaphase, not later, but remaining attached to the telomeres until telophase. Were tethers to produce backwards forces at late anaphase, when the poleward forces cease, they could move the late anaphase chromosomes backward, negating the proper anaphase distribution of chromosomes into new cells.

In the present experiments we wanted to establish a partial lysis system in order to study tethers without the need for a laser microbeam and under conditions in which one might be able to add normally impermeable components to the cell in order to probe the tethers. To explain our tactic for attaining this goal we first need to explain one more property of tethers, that elastic tethers become inelastic because of dephosphorylation by protein phosphatase 1 (PP1).

Calyculin A (CalA), an inhibitor of serine-threonine protein phosphatase 1 (PP1) and of protein phosphatase 2A (PP2A), when added to early anaphase cells causes chromosomes to move backwards at the end of anaphase, telomere moving to partner telomere. Fabian *et al*. (2007a), who originally described that result, suggested that the backwards movements were due to mitotic tethers, and that CalA treatment (blocking dephosphorylation) caused the tethers to remain elastic. Further indication that this is so is that when CalA was added at different times after the start of anaphase there was less and less backward movement of the chromosomes when the CalA was added later and later in anaphase (Kite and Forer, 2020); this matches very well the behavior of arm fragments produced in control cells where there is less arm fragment movement when arms are produced when tethers are longer than when arm fragments are produced when tethers are shorter. Because CalA inhibits PP1 and PP2A equally well, but okadaic acid at the same concentrations inhibits only PP2A, Fabian et al. (2007b) and Kite and Forer (2020) treated cells with okadaic acid: the chromosomes did not move backwards. Thus, the dephosphorylation events causing backwards movements are due to PP1. From their experiments treating cells with CalA, Kite and Forer (2020) posited that at the beginning of anaphase tethers most likely are phosphorylated and elastic; that as anaphase progresses the tethers gradually are dephosphorylated; and that at end of anaphase tethers are completely dephosphorylated and inelastic. Subsequent experiments utilizing laser microbeam irradiations verified that the CalA-induced backwards movements indeed are due to tethers (Forer et al., 2021): after cells were treated with CalA, cutting either telomere stopped the backwards movements of the chromosomes; severing a leading arm stopped the backwards chromosome movements as the arm fragment continued to move backward; and directly severing tethers stopped chromosome backward movements.

The present experiments are aimed at obtaining functional tethers in partially lysed cells. We first tested if the tether properties we just described have been altered or not by the lysis buffer. We aimed for partially lysing the cells, to make them permeable, and at the same time we needed to inhibit the force production that moves the chromosomes poleward. The prediction is that if the poleward forces are abolished but the backwards forces from the tethers are still active, then the tethers will function to move the anaphase chromosomes backwards when the tethers are elastic (i.e., short tethers, early anaphase), but will not move the chromosomes backwards when they are not elastic (i.e., long tethers, late anaphase). A second prediction is that if we partially lyse cells treated with CalA, the chromosomes should move backwards even if the tethers are long when we permeabilize the cells. To further ensure that the tether properties are not changed by the lysis medium we also measured other properties. For one, there is extensive data for tether-length dependence of arm fragment movements as the fractional distance (fraction of tether length) the arm fragments move when arm fragments are formed at different tether lengths (Forer et al., 2021): we measured that in the partial-lysis system. We also compared velocities: although shorter tethers are more elastic than longer tethers, they do not cause faster backward movements than longer tethers (Forer et al., 2021). Furthermore, treating cells with CalA in early anaphase maintains tether elasticity until the end of anaphase, but the maintained elasticity does not increase the velocities of arm fragments (Forer et al., 2021); thus, in the partial lysis system we measured the velocities of chromosomes after cells were treated with CalA.

Our objective is to test whether elastic tethers continue working to cause backward chromosomal movements in a normal way despite the arrested anaphase spindle apparatus that no longer moves chromosomes poleward. If tethers function normally in the partially lysed *in vitro* system then the system could be developed further, for example, to being able to add phosphorylation promoters/inhibitors directly to permeabilized cells and analyzing their effect on tether elasticity. Having an accessible system to study tethers means that tethers can be studied at any point during anaphase using protocols which do not depend on laser cutting of chromosomal arms.

In the research presented herein, a non-invasive procedure was devised in which anaphase spindle poleward forces were deactivated by partially lysing the cell; tether function was then assessed and found to be the same as in non-treated cells.

## MATERIALS AND METHODS

### Living cell preparation

Crane flies (*Nephrotoma suturalis* Loew) were reared in the lab using procedures similar to those described by Forer (1982). Procedures to make living cell preparations are described in detail by Forer and Pickett-Heaps (1998). Briefly, testes from IV-instar larvae were dissected under halocarbon oil, placed into a drop of halocarbon oil, then washed thrice in insect Ringer’s solution modified (by adding a buffer) from that described by Ephrussi and Beadle (1936), (0.13 M NaCl, 5mM KCl, 1.5 mM CaCl_2_, 3mM phosphate buffer, pH 6.8). A testis then was placed in a 2.5 µL drop of insect Ringers solution containing fibrinogen (10mg/mL) on a coverslip, and was pierced to spread out the cells. These cells then were fixed in place by adding 2.5 µL thrombin to the fibrinogen to form a fibrin clot (Forer and Pickett-Heaps, 2005). The coverslip with cells embedded in the fibrin clot was inverted over insect Ringers solution in a perfusion chamber (Forer and Pickett-Heaps, 2005), and sealed with a molten mixture of 1:1:1 vaseline, lanolin, and paraffin. Ringer solution then was perfused through the chamber and the perfusion chamber was placed on the stage of a phase-contrast microscope for observation.

### Different cell treatments

The dividing cells in the perfusion chamber were studied using the phase-contrast microscope and images recorded in real time using a DVD recorder. Once the cells were at the required stage of division they were perfused according to the treatment group they belonged to. Control cells were perfused with Ringers solution and the experimental cells were perfused with dilutions of the lysis buffer we use for immunofluorescence staining (e.g., Fabian et al., 2007b; Sheykhani et al., 2017). The lysis buffer (100 mM piperazine-N, N-bis (2-ethanesulfonic acid) [PIPES]; 10 mM EGTA; 5 mM MgSO_4_; 5% DMSO; 1% IGEPAL CA-630; pH 6.9), essentially a cytoskeleton stabilizing buffer to which detergent is added, was diluted in Ringers solution by (1:1000), (1.5:1000), (2:1000), (2.5:1000), (4:1000), (5:1000), or (6:1000). Each lysis buffer dilution was added to the cells separately; 10-15 minutes after lysis buffer treatment, some cells were perfused with Ringers solution to wash out the lysis buffer to test whether the effects of the lysis buffer were reversible. For experiments using Calyculin A (CalA) (LC Laboratories, Woburn, MA), 50µM CalA was prepared in DMSO and stored as frozen aliquots. For experimentation, aliquots were thawed and diluted 1:1000 in Ringers solution. The resultant 50nM CalA was added to the cell in early anaphase. Later in anaphase we added to the cells 50nM CalA diluted 1:1000 from 50µM CalA with diluted lysis buffer (lysis +CalA) at the lysis buffer concentrations used for the initial perfusions. For experiments in which we tried to identify which components of the lysis buffer were important, individual lysis buffer components were prepared separately in Ringers solution at their final concentration found in 6:1000 lysis buffer, namely: 0.56% PEM (PIPES, EGTA and MgSO_4_ combination), 0.006% IGEPAL, and 0.03% DMSO. The final working concentration of DMSO in the 6:1000 lysis experiments was 0.03%, but when testing the individual components DMSO was used at 1% to verify that up to 1% DMSO in experimental solutions have no effect on the experimental results (LaFountain, 1985; Silverman-Gavrila and Forer, 2000). Some of the 6:1000-lysis-buffer-treated cells were prepared for immunofluorescence by lysing them with full-strength lysis buffer after 25-35 minutes of treatment with 6:1000 lysis buffer.

### Microscopy and data analysis

The living cell preparations were studied using phase-contrast microscopy with a *Nikon* 100X, 1.25 NA phase-contrast oil immersion objective lens. Real-time video images were recorded on DVDs which were later converted into time-lapse video sequences (.avi files) using freeware *VirtualDub 2*. Individual video-sequence frames were analyzed; measurements of chromosomal movements were made using an in-house program *WinImage* (Wong and Forer, 2003). Chromosome-movement graphs were plotted using the commercial software *SlideWrite,* version *Plus 7.0* (Forer and Berns, 2020).

### Fluorescent staining and confocal microscopy

We followed the immunostaining procedure of Fabian and Forer (2005). Control and experimental cells were lysed in full-strength lysis buffer for 15 minutes. The cells then were fixed in 0.25% glutaraldehyde in phosphate-buffered saline (PBS) for 2-3 minutes, rinsed twice in PBS (5 minutes each rinse), placed in 0.05M glycine for 10 minutes (to neutralize the free aldehyde groups), and rinsed four times in PBS (5 minutes each rinse). Finally, the coverslips with the fixed cells were stored in PBS-glycerol 1:1 (v/v) mixture at 4°C. For immunostaining, the stored coverslips were first washed in PBS to remove all the PBS-glycerol mixture, rinsed with 0.1% Triton X-100 in PBS to ensure the antibodies spread evenly over the cell preparation, and then treated sequentially with two primary antibodies. They were stained first for tyrosinated α-tubulin and then for acetylated α-tubulin. The antibody to tyrosinated α-tubulin was added first using rat monoclonal antibody YL1/2 (Abcam), diluted 1:200 with PBS. This was followed by staining with mouse-absorbed Alexa 488-conjugated goat anti-rat antibody (Molecular Probes) diluted 1:50 with PBS. Acetylated α-tubulin antibody was added next, using as primary antibody mouse monoclonal antibody 6-11B-1 (Millipore Sigma) diluted 1:50 with PBS. This was followed by staining with rat-absorbed Alexa 568-conjugated donkey anti-mouse antibody (Invitrogen) diluted 1:200 with PBS. All cells were incubated in each antibody for 1 hour and kept in the dark to prevent fluorochrome inactivation by light. After each antibody incubation, cells were rinsed twice in PBS (5 minutes each rinse), then rinsed with 0.1% Triton X-100 in PBS to ensure the next antibody spread evenly over the cells. After the last antibody staining, coverslips were rinsed twice in PBS (5 minutes each rinse), and then placed in PBS-glycerol 1:1 (v/v) for 2-3 minutes to prepare for mounting. Finally, coverslips were mounted on slides in Mowiol solution (Osborn and Weber, 1982) to which we added paraphenylene diamine as antifading agent (Platt and Michael 1983), and left to dry in the dark for 24-48 hours. Once dry, slides with the stained cells were stored in the dark at 4°C. When ready to analyze, cells were studied using an LSM 700 Zeiss Observer confocal microscope, with a Zeiss Plan-Apochromat 63X 1.4 NA oil-immersion objective lens. Images were collected using ZEN Black software, and further processed using FIJI Image J software.

## RESULTS

### Control cells

Primary spermatocytes of crane flies have three bivalent autosomes and two unpaired sex chromosome univalents. During normal meiosis-I (Figure 1), the bivalent autosomes and the univalent sex chromosomes line up at the equator in metaphase. Anaphase begins (Figure 1, b) as the three bivalents disjoin into six half-bivalents which then move towards their respective poles. As the autosomes move poleward the spindle length remains constant (Forer, 1966) and both sex chromosomes remain stationary at the equator (Forer *et al.,* 2013). The half-bivalents approach their respective poles in about 20-30 minutes after which the spindle starts elongating, the sex chromosomes start segregating (Figure 1, g, h, i), and the cleavage furrow ingresses.

**Figure 1.**
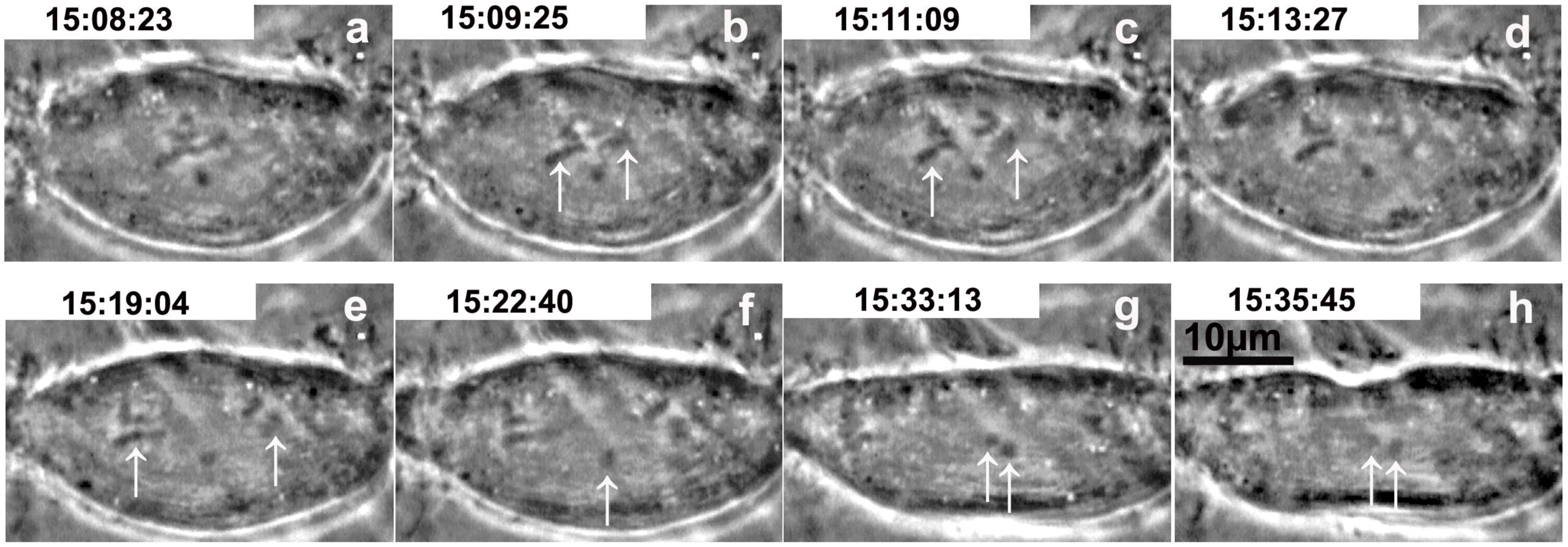
Meiosis-I in a non-treated (control) crane-fly primary spermatocyte. Time (hrs:min:sec) is at the top of each image panel. **(a)** Metaphase bivalents. **(b–e)** Arrows indicate the positions of two separating half-bivalents at the start of anaphase **(b)** and as they move in opposite directions to their respective poles **(b-e)**. Arrows in **(f-h)** indicate the position of the two sex chromosomes; at first both are at the equator **(f)** and then they move towards opposite poles **(g-h)** as the autosomes get near the poles. The spindle elongates as both sex chromosomes separate, and the cleavage furrow appears **(h).** Scale bar in **(h)** represents **10 µm.**

### Experimental cells

#### (1) Dilute lysis buffer treatment stops anaphase chromosome movements; chromosomes then move backward

Cells at different stages of anaphase were treated with diluted lysis buffer. That treatment often stopped anaphase chromosome movement. As an example, Figure 2A shows a cell before and after treatment with lysis buffer (diluted 6:1000 with Ringers solution). Two chromosomal pairs were in the plane of focus (Figure 2A *a* and *b*, arrows) and both pairs stopped moving poleward and began to move backward (arrows in *d, e* and *f*). We determined the effects on chromosome movements by plotting the positions of telomeres and kinetochores before and after addition of diluted lysis buffer. Figure 2B shows the resultant graph of positions *versus* time. For measurements, a fixed point was chosen near the right spindle pole (arrowhead in Figure 2A, *a*). from which the distances of kinetochores and telomeres for each chromosomal pair were measured. As seen in Figure 2B, immediately after lysis treatment the distance between partner telomeres was 3 µm, which is the initial tether length. Both telomeres and kinetochores stopped moving apart (normal chromosome movement stopped) and almost immediately the kinetochores and telomeres moved towards each other (the partner chromosomes moved backward). At the end of their backward movement the telomeres were touching which means that the tethers connecting the telomeres shortened by 100% and thus the *fractional distance* (of the initial tether length) that encompassed chromosome backward movement was 1. Figures 3A and 3B illustrate another cell, this one treated with lysis buffer diluted 1:1000. In the one chromosome pair we were able to follow, the lysis buffer almost immediately stopped the poleward chromosome movement and almost immediately thereafter the two half-bivalents started to move backward. The telomeres moved backward until they eventually touched each other (Figure 3B).

**Figure 2A.**
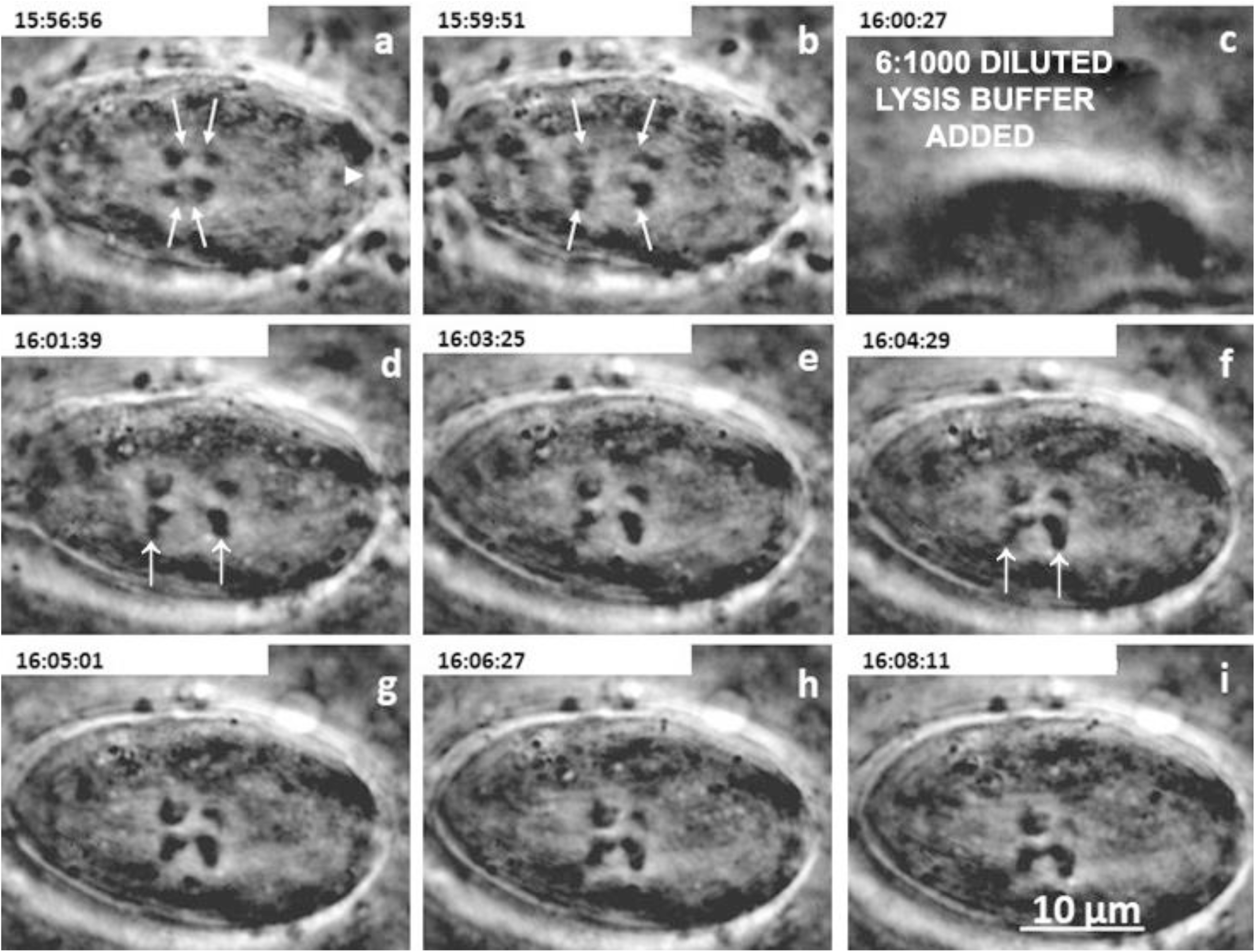
A cell treated with 6:1000 lysis buffer. Time (hr:min:sec) is at the top of each panel. **(a)** Two half-bivalent pairs start anaphase; each of the visible 4 half-bivalents is indicated by arrows. An arrowhead indicates the point near the right spindle pole used as a fixed reference point for measurements. **(b)** Partner half-bivalents continue separating. **(c)** Dilute lysis buffer added. **(d-g)** The half-bivalents (arrows) start moving backward towards each other. **(h)** Backward movements stop as partner telomeres meet each other. **(i)** Partners remain as they are. Scale bar represents **10 µm.**

**Figure 2B.**
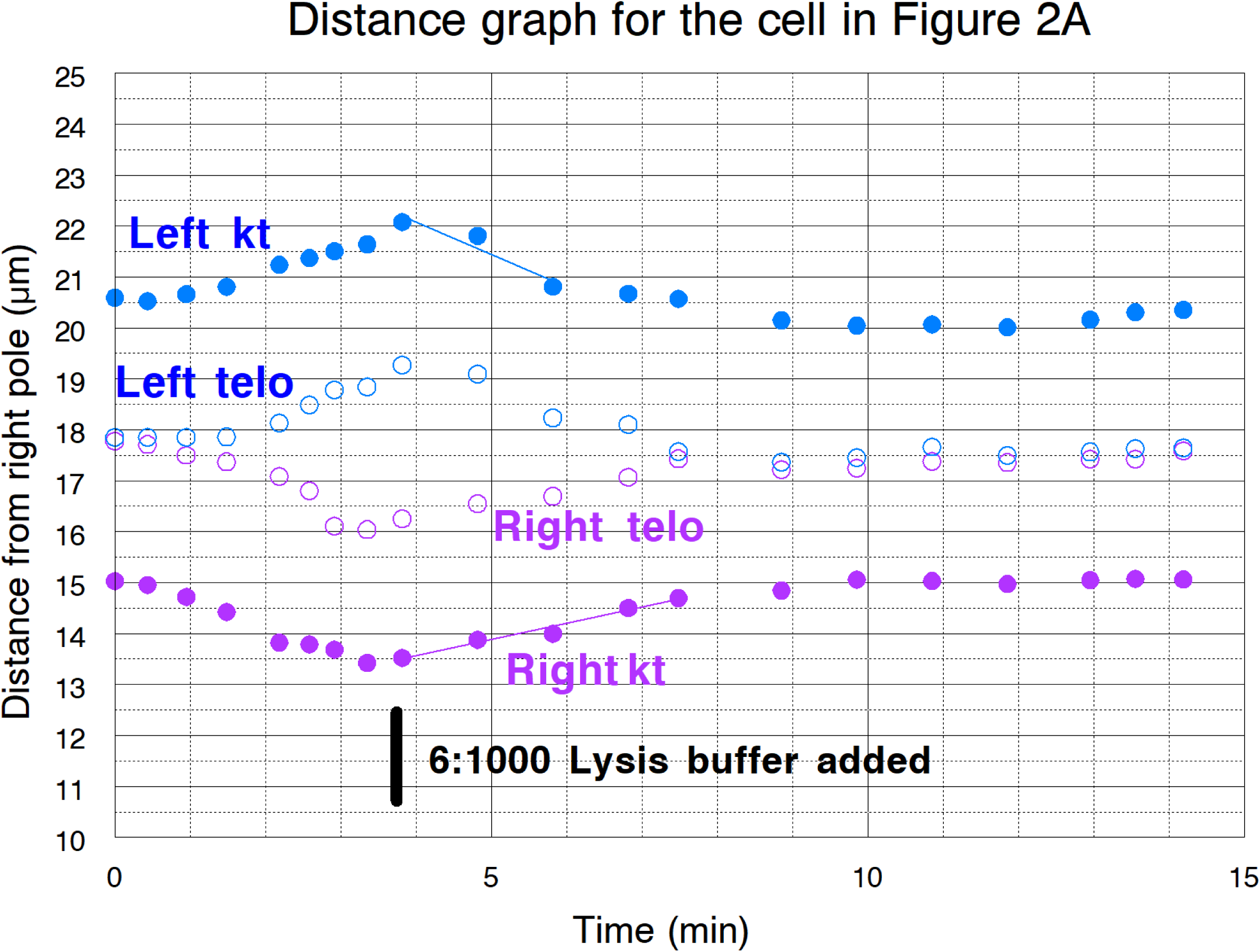
Movement graph for the bottom half-bivalent pair illustrated in Figure 2A. The graph time of 0 min corresponds to Figure 2A panel (a), image time=15:56:56 (hr:min:sec). Distance measurements were made from a point near the right pole (Figure 2A, a). Partner kinetochores are marked as *Right kt* and *Left kt*, partner telomeres are *Right telo* and *Left telo*. Partners were moving apart when 6:1000 lysis buffer was added. After lysis buffer addition they stopped moving poleward and moved backwards until the telomeres touched each other. Slopes of the individual graphs represent movement velocities.

**Figure 3A.**
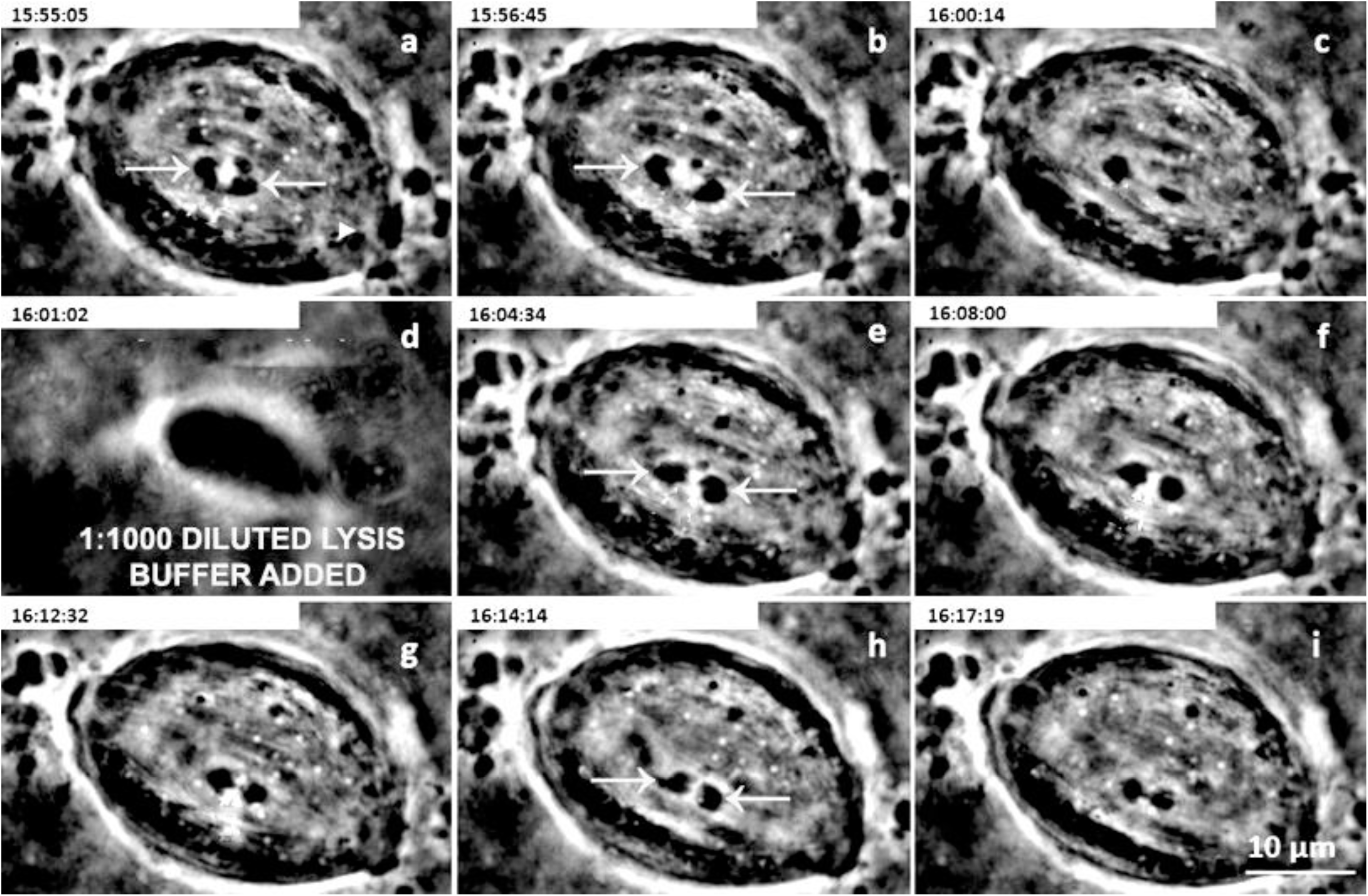
A cell treated with 1:1000 lysis buffer. Times (hr:min:sec) are indicated at the top of each panel). **(a)** Bivalents start anaphase separation. Two partner half-bivalents are indicated by arrows. A single arrowhead indicates the point near the right spindle pole used as a fixed reference point for measurements. **(b-c)** Partner half-bivalents continue separating. The same two as in **a** are indicated by arrows (as well as in **e** and **h**). **(d)** Lysis buffer added. **(e-g).** Partners stop separating and start moving backward toward each other. **(h)** Backward movements stop as partner telomeres meet. **(h-i)** Partners remain as they are. Scale bar in **(i)** represents **10 µm.**

**Figure 3B.**
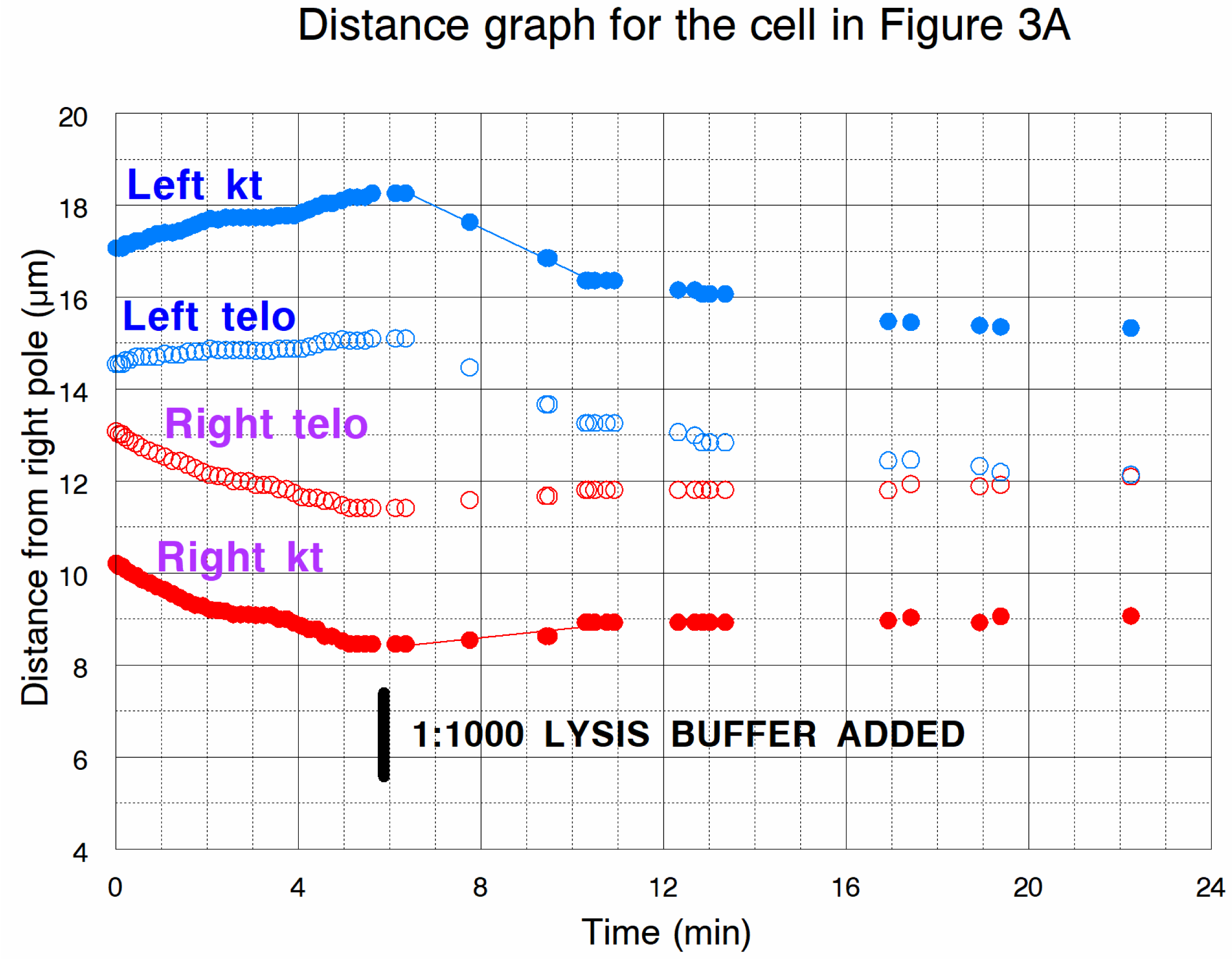
Distance *versus* time graph for the partner half-bivalents illustrated in Figure 3A. The graph time of 0 min corresponds to image time=15:55:05 in (Fig. 3Aa). Distance measurements were made from a fixed point near the right pole [arrowhead in Fig. 3A**(a)**]. Partner kinetochores are marked as Right kt and Left kt, partner telomeres are Right telo and Left telo. Partner telomeres were moving apart when lysis buffer was added after which they stopped separating, moved backward, and met each other. Slopes of the lines on the graphs represent movement velocities.

Not all chromosome pairs in cells treated with 1:1000 lysis buffer stopped their poleward anaphase movement. One such cell is illustrated in Figures 4A and 4B. The images are of one half-bivalent pair (though we have data from all three pairs), one of the three pairs in the cell in which all partners continued their anaphase movements towards the poles, albeit at reduced speeds (Figure 4B).

**Figure 4A.**
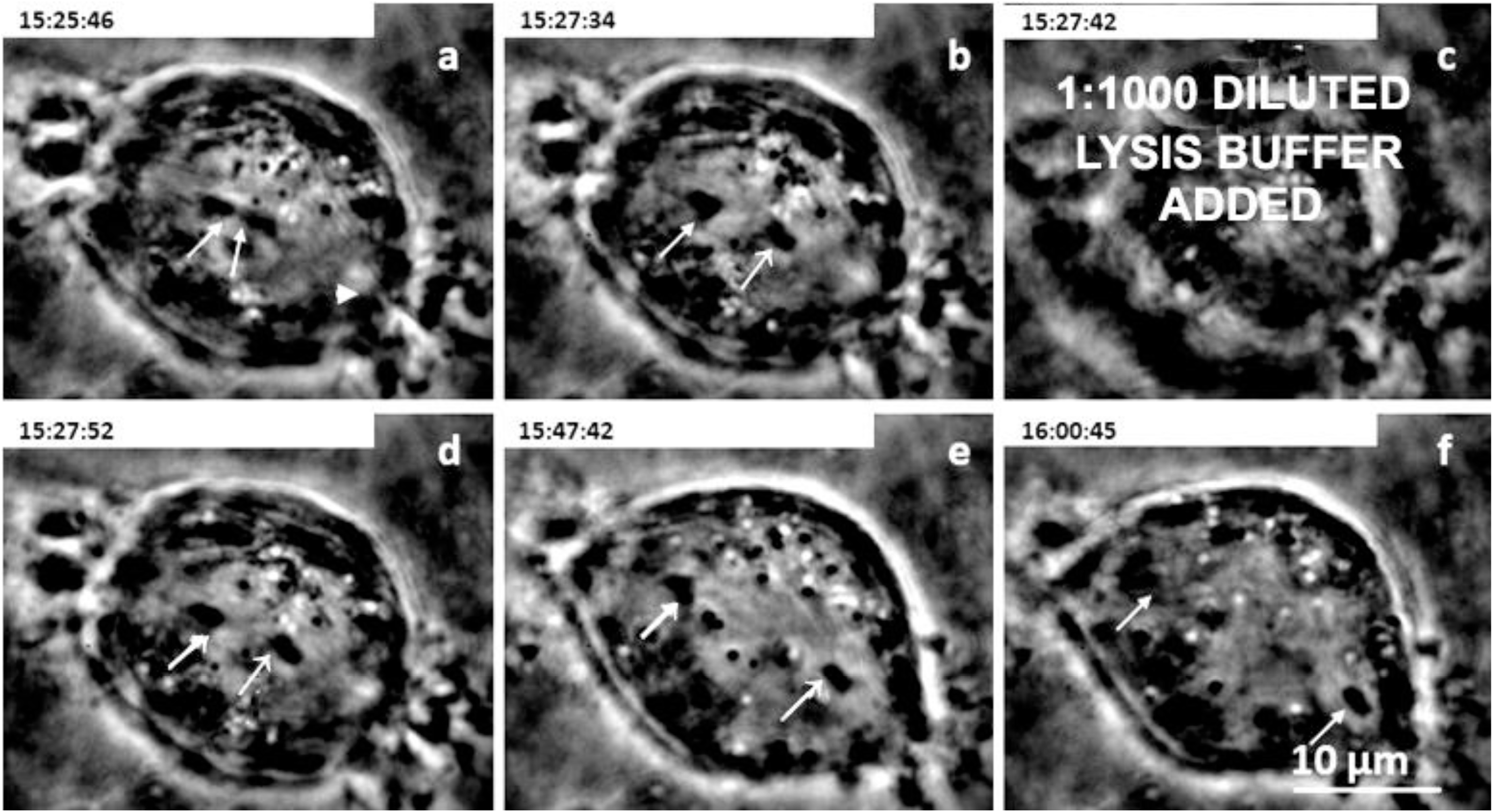
A **c**ell treated with 1:1000 lysis buffer. Times (hr:min:sec) are indicated at the top of each panel. **(a)** Half-bivalents start anaphase separation; the same set of partner half-bivalents is marked by arrows in all frames. A single arrowhead in (**a)** indicates a point near the right spindle pole used as a fixed reference point for measurements. **(c)** Lysis buffer added. **(d)** Poleward movements slow-down. **(e)** poleward movements continue at reduced speeds. **(f)** Partner half-bivalents are near to their poles. Scale bar in **(f)** represents **10 µm.**

**Figure 4B.**
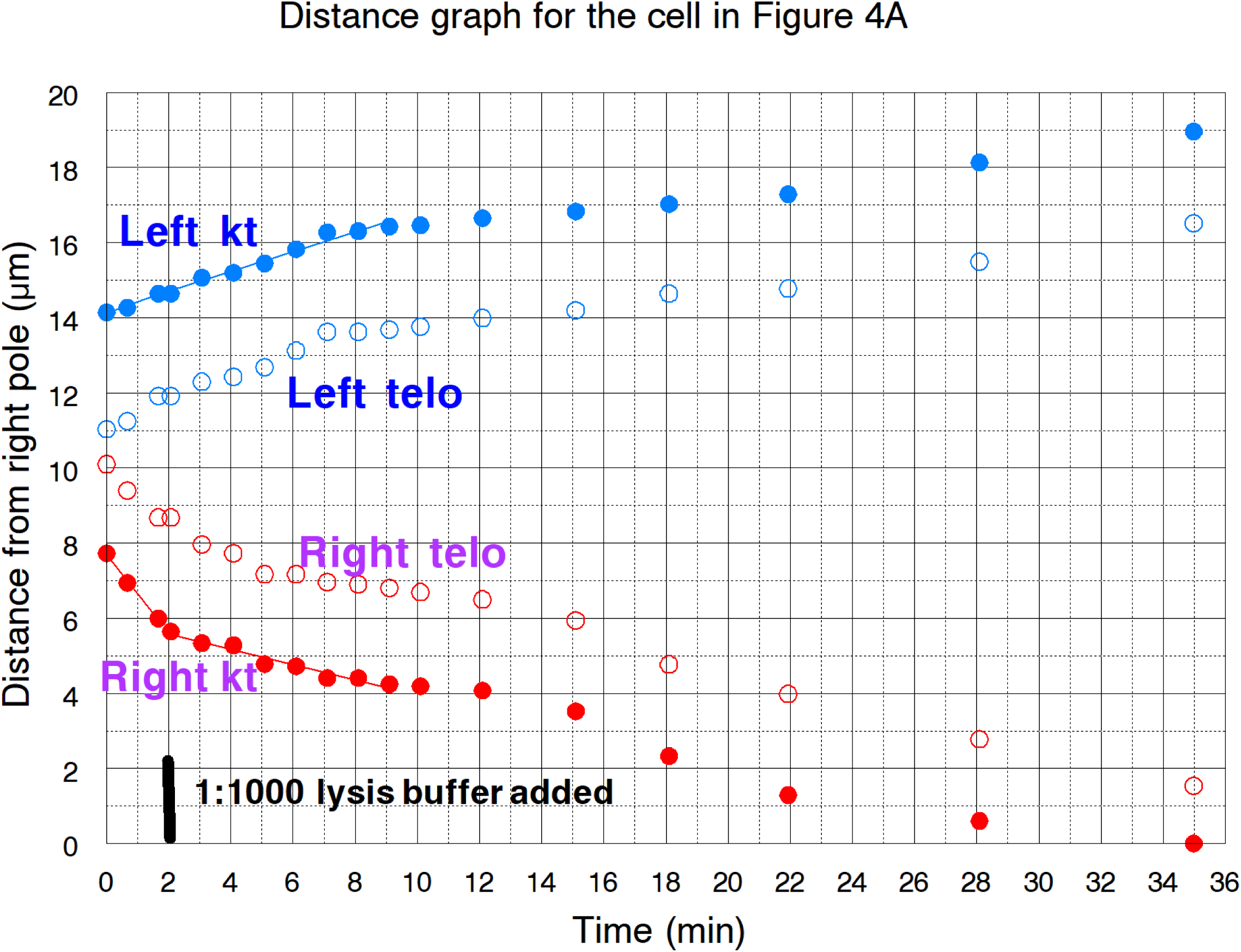
Distance *versus* time graph for the half-bivalent pair illustrated in Figure 4A. The graph time of 0 min corresponds to the Figure 4A(a) image at 15:25:46 (hr:min:sec). Distance measurements were made from a point near the right pole (arrowhead in 4Aa). Partner kinetochores are marked as Right kt and Left kt, partner telomeres are Right telo and Left telo. Partners were moving apart when lysis buffer was added, after which one half-bivalent immediately slowed down, both half-bivalents continued to move poleward, and later the other half-bivalent slowed down.

We treated some cells with Calyculin A (CalA) before adding lysis buffer. CalA-pretreated cells seem to respond differently from control (not Cal-treated) cells after treatment with lower concentrations of dilute lysis-buffer in that anaphase always stopped after treatment with diluted lysis buffer (Table 1, Figure 5).

**TABLE 1.** Effects of different concentrations of lysis buffer on poleward anaphase chromosome movements.

| Lysis buffer<br>dilution (μL per<br>1000μL Ringers) | Anaphase poleward<br>movement **stopped** |  | Anaphase poleward<br>movement **slowed** |  |
| --- | --- | --- | --- | --- |
|  | Treated cells | Half-bivalent pairs<br>Stopped/observed | Treated cells | Half-bivalent pairs<br>Slowed/observed |
| <3 | 13 | 36/36 | 6 | 15/15 |
| 4 – 5 | 4 | 10/10 | 0 | 0 |
| 6 | 20 | 55/55 | 0 | 0 |
| 50nM CalA followed<br>by 50nM CalA in lysis<br>buffer (μL per1000μL<br>Ringers) |  |  |  |  |
| <3 | 4 | 9/9 | 0 | 0 |
| 6 | 3 | 8/8 | 0 | 0 |

**Figure 5.**
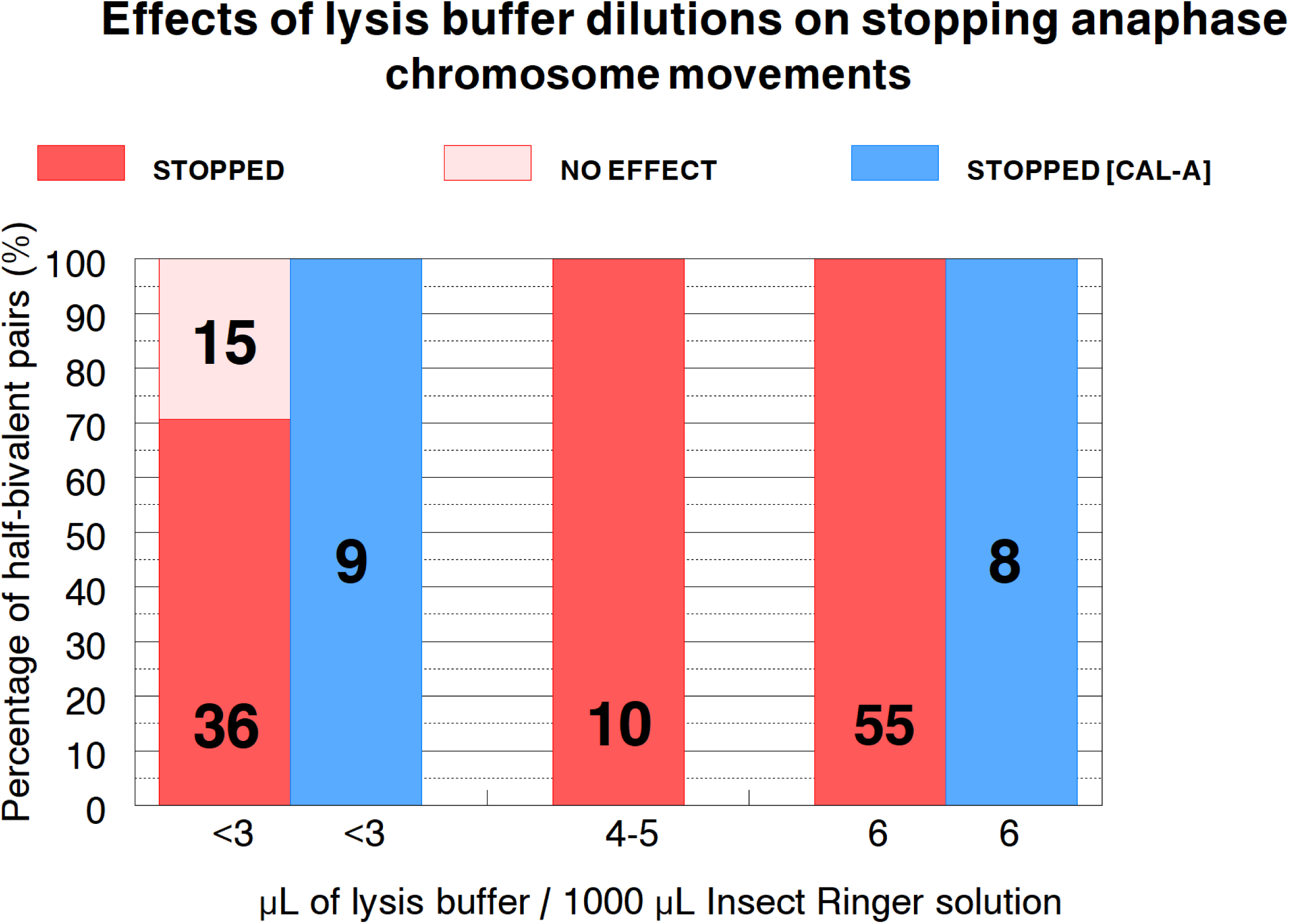
**Graphical representation of the data in Table 1** showing effects of lysis buffer treatments on stopping anaphase chromosome movements in both normal cells and in cells pre-treated with CalA. Each bar represents the percentage of chromosome pairs that stopped/did not stop poleward movement at the lysis buffer concentrations indicated on the abscissa. The blue bars represent cells pre-treated with CalA. 133 half-bivalent pairs were examined in total; the numbers of half-bivalent pairs assayed with each lysis concentration are written on the respective bars.

We aggregated the data from the different concentrations of diluted lysis buffer. We first consider, in aggregate, the effects of the dilute lysis buffers on poleward chromosome movements during anaphase. Then we consider the effects on backward movements of the chromosomes.

With regard to effects on anaphase chromosome movement, in every cell measured, each of the three separating half-bivalent pairs reacted the same: all pairs either stopped poleward movements or slowed them down (Table 1). Thus the buffer caused changes to the entire spindle apparatus so that poleward forces either were significantly reduced or were brought to zero. That some of the chromosome pairs slowed but not stopped (Table 1, Figure 5), shows that when poleward movement was not stopped completely, the treatments nonetheless interfered with normal poleward force production.

A second conclusion is that dilute lysis buffer consistently stops poleward anaphase chromosome movement only when concentrations of dilute lysis buffer are above 3µL/1000µL: chromosome movements sometimes stop when lysis-buffer concentrations are less than 3µL/1000 µL, but not as consistently as when using higher concentrations (Table 1, Figure 5). Backward movement occurred only when poleward movement was stopped, not when it was slowed; this presumably is because the force for the slowed poleward movement was still greater than the backward force from the tethers.

#### (2) Cells in which chromosomes stopped moving after treatment with dilute lysis buffer can return to normal after rinsing with Ringer solution

We tested whether diluted lysis-buffer kills the treated cells when it stops anaphase chromosome movements. We washed out the diluted lysis-buffer with Ringers solution between 10-15 minutes after the initial application of the diluted lysis buffer. When cells recovered, all half bivalents in each cell resumed segregation within 5-15 minutes of washout: the average recovery time before chromosomes resumed poleward movement was 8 minutes post-wash. After washout, those cells that recovered were normal – that is, chromosomes moved to poles and telophase and cleavage ensued. All half-bivalent pairs in each cell acted the same, but not all cells recovered from lysis-buffer treatment. When the concentration of the diluted lysis-buffer reached 6µL of lysis buffer/1000µL Ringers, the less likely that the cells recover (Figure 6, Table 2). Thus, the high concentration of lysis buffer seem to cause irreversible, deleterious changes to the cells.

**Figure 6.**
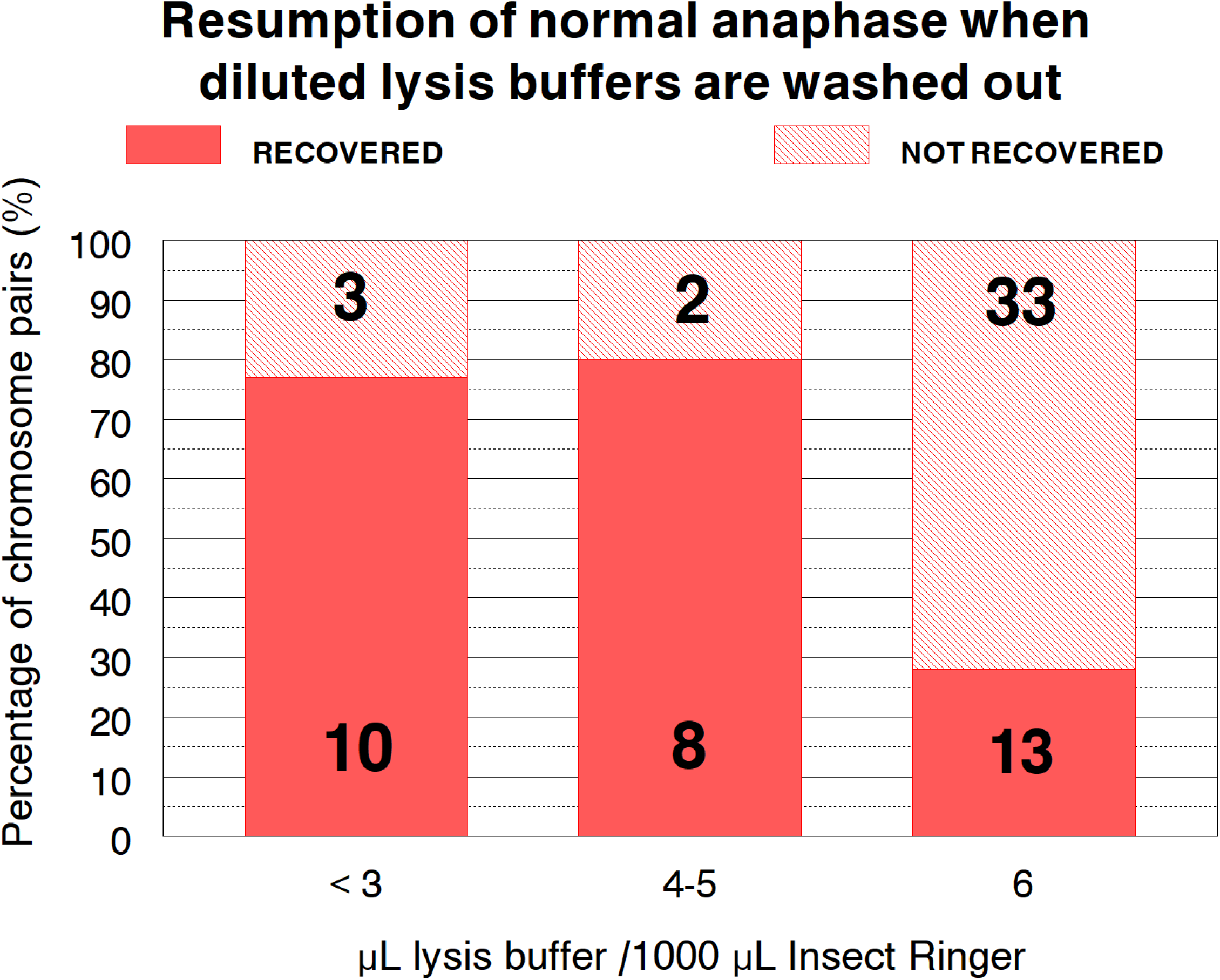
Graphical representation of the data in Table 2 on recovery of normal anaphase after washing out the dilute lysis buffer. We measured whether anaphase movement recovered after washing out the dilute lysis buffers (concentrations listed on the abscissa) with Ringers solution. The numbers on each bar represent the numbers of half-bivalent pairs with the specified result at the specified concentration of dilute lysis buffer. The total number examined was 69 half-bivalent pairs.

**TABLE 2.** Recovery of anaphase movement after washing out dilute lysis buffer with Ringer solution.

| Lysis buffer dilution<br>($\mu\text{L}$ buffer/1000 $\mu\text{L}$<br>Ringers) | Chromosome movement<br>**recovered** after washout | | Chromosome movement **not**<br>**recovered** after washout | |
| --- | --- | --- | --- | --- |
|  | Cells | Pairs recovered/pairs<br>observed | Cells | Pairs not recovered/<br>pairs observed |
| < 3 | 4 | 10/10 | 1 | 3/3 |
| 4 – 5 | 3 | 8/8 | 1 | 2/2 |
| 6 | 5 | 13/13 | 12 | 33/33 |

We now describe the backward movements that occur after the diluted lysis buffer stopped normal anaphase chromosome movements to the pole.

#### (3) After treatment with diluted lysis buffer, shorter tethers move chromosomes backward more consistently than longer tethers do

We determined whether backward chromosome movements in cells treated with dilute-lysis buffer varied with tether length (at the time the lysis buffer was added). There were no obvious differences in backward movement with different concentrations of dilute-lysis buffer, including treatments with high concentrations that were not reversible. The only requirement to have chromosomes move backward was that the treatment stop poleward anaphase movement. Thus, we analyzed the aggregated data, the complete set of data listed in Table 1, without regard to concentration of diluted lysis buffer.

Backward movements started within 1 minute of anaphase arrest. [We considered any chromosome movements as “backward” when the tether length decreased by >0.5µm.] As illustrated in Figure 7 (pink bars), tethers that were shorter at the time of lysis-buffer addition were more likely to move chromosomes backward than were tethers that were longer. In Cal-A pre-treated cells chromosomes moved backward all the time, at all tether lengths, even in cells that had very long tethers at the time of lysis-buffer treatment (Figure 7, blue bars). To test whether this is a real difference, that is, whether the frequencies of backward movement in cells pre-treated with CalA are statistically different from the values in non-pre-treated cells, we used (33/62) as the probability for getting backward movement at tether lengths at 5 µm or above for not-pre-treated cells shown in Figure 7. We asked what the probability would be for 17 pairs in a row to move backward with that probability for each one (we used 17 because all 17/17 half-bivalent pairs moved backward after pre-treatment with CalA). The chance of getting 17 in a row to move backward, each with a probability of 33/62, is (33/62)^17^, which is ∼ 2x 10^-5^. This very low probability indicates that the backward movement in CalA treated cells is not a function of tether lengths, that CalA prevents loss of backward chromosome movement as tethers are longer. Thus movements of half-bivalents in the cells treated with dilute lysis buffer are the same as movements of laser-induced arm fragments (Forer et al., 2021): movements of arm fragments in non-treated cells depends on lengths of the tethers when the arm fragments are formed but movements of arm fragments in cells treated with CalA (in early anaphase) do not depend on lengths of tethers: chromosome arm fragments moved to their partner telomeres at all tether lengths, even very long tether lengths in which arm fragments in control cells never moved backward. This shows that backward movements in our model lysis system is the same as tether-generated movements of arm fragments used to identify forces from mitotic tethers between partner telomeres.

**Figure 7.**
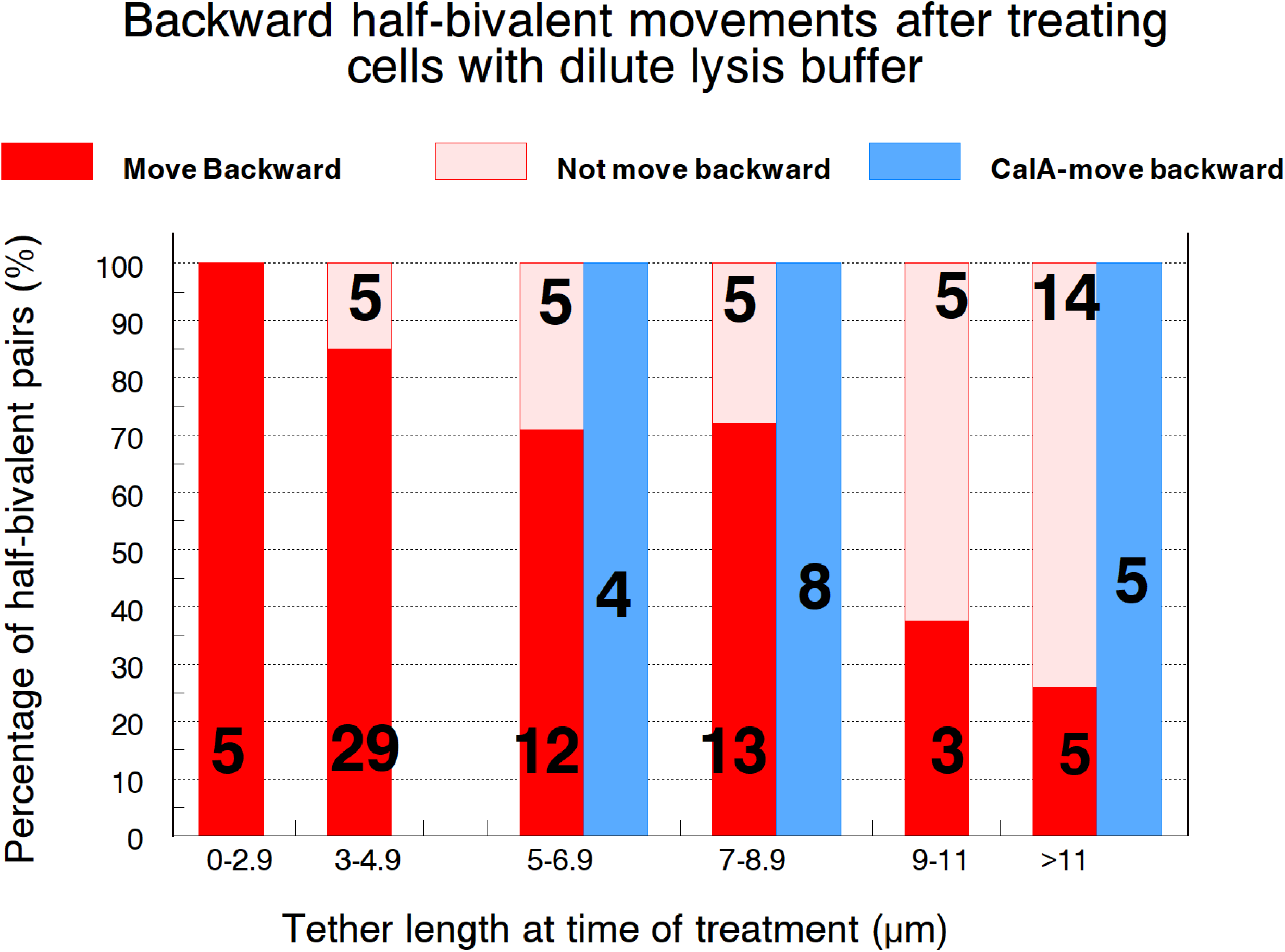
Frequencies of backward movement [of **≥0.5 µm]** with different initial tether lengths for cells treated with lysis buffer (pink/red bars) and for cells pre-treated with CalA before adding lysis buffer (blue bars). 118 half-bivalent pairs were examined; numbers of each tether-length category are written on the respective bars. Lysis buffer dilutions were between <3 to 6 µL per 1000µL Ringers solution. Differences between lysis only and CalA pre-treated cells are statistically significant since statistical treatment (t-tests) indicate that the probability of them being due to chance is <0.01.

#### (4) After dilute lysis buffer treatment, chromosomes with shorter tethers move backward for longer fractional distances of the initial tether lengths than chromosomes with longer tethers do

The distance moved backward by partner half-bivalents when taken as a fraction of the initial tether length reflects how much the elastic tether shortens as it pulls its partner half-bivalent backward. With respect to chromosome arm fragments, the distance that arm fragments move (as a fraction of initial tether length) decreases as tether lengths increase (e.g., Figure 9B of Forer et al., 2021). With respect to chromosomes in cells treated with dilute lysis buffer, we measured the fractional shortening of tethers during backward chromosome movement (Figure 8A). To determine if there is a statistically significant correlation between tether length and fractional distance moved backward with lysis treatment alone, we statistically analyzed the data, omitting one data point from Figure 8A at tether length of 8.6 µm because it had fractional backward movement of less than 0.10; we did this because such small transient movement might not have been due to tether elasticity but rather due to partner homologues briefly propelling backwards as poleward-directed anaphase forces were deactivated. The two variables that were analyzed were the fractional distances of initial tether lengths that the chromosomes moved backwards, and the tether lengths at the time of anaphase arrest. Since the two were non-parametric, Spearman’s rank correlation was computed to assess their relationship (Figure 8B). Spearman’s correlation coefficient (r_s_) was -0.610, and with P ≤ 0.01, the critical value for r_s_ at the given sample size of 100 was 0.257. Therefore, during lysis treatment, fractional distance moved backward by chromosomes due to their tether elasticity is negatively correlated to the tether length more than 99% of the time (Figures 8A, 8B). Hence, the backward movement caused by tethers as a fraction of initial tether length varies with initial tether length: at shorter tether lengths the chromosomes move larger fractions of the initial tether lengths than at longer tether lengths. This relationship, illustrated in Figure 8C, is similar to backwards movement of arm fragments (Figure 9 of Forer et al., 2021), and is yet further corroboration that tether elasticity and functioning is unaffected by the treatments with dilute lysis buffer.

**Figure 8A.**
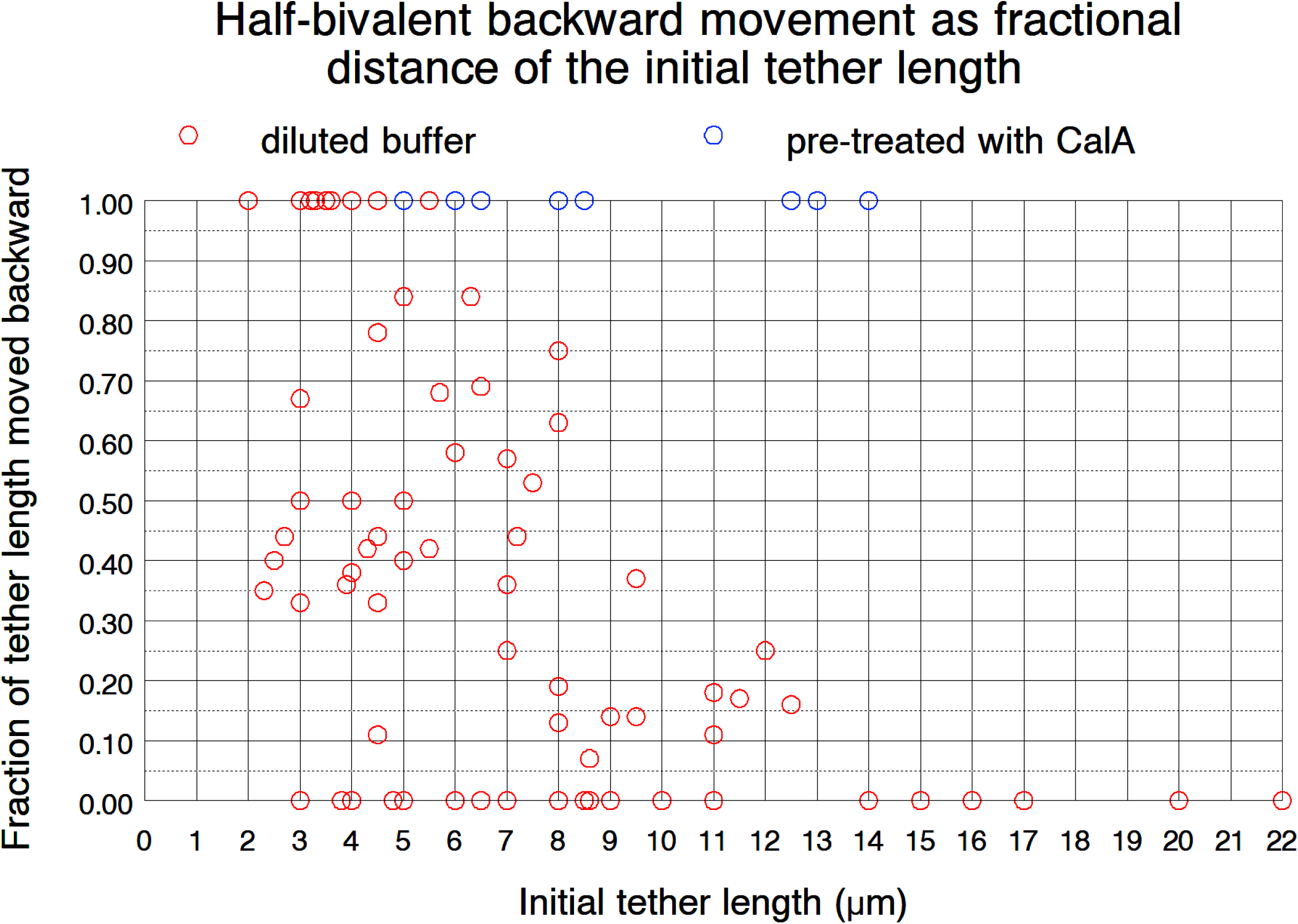
Distances half-bivalents moved backwards as fractions of their initial tether lengths. Cells were treated with lysis buffer (red circles) or were pre-treated with CalA before lysis (blue circles). 118 pairs of half-bivalents were examined in total. The data points are in aggregate, so different points on the graph are not necessarily treated with the same dilution of lysis buffer.

**Figure 8B.**
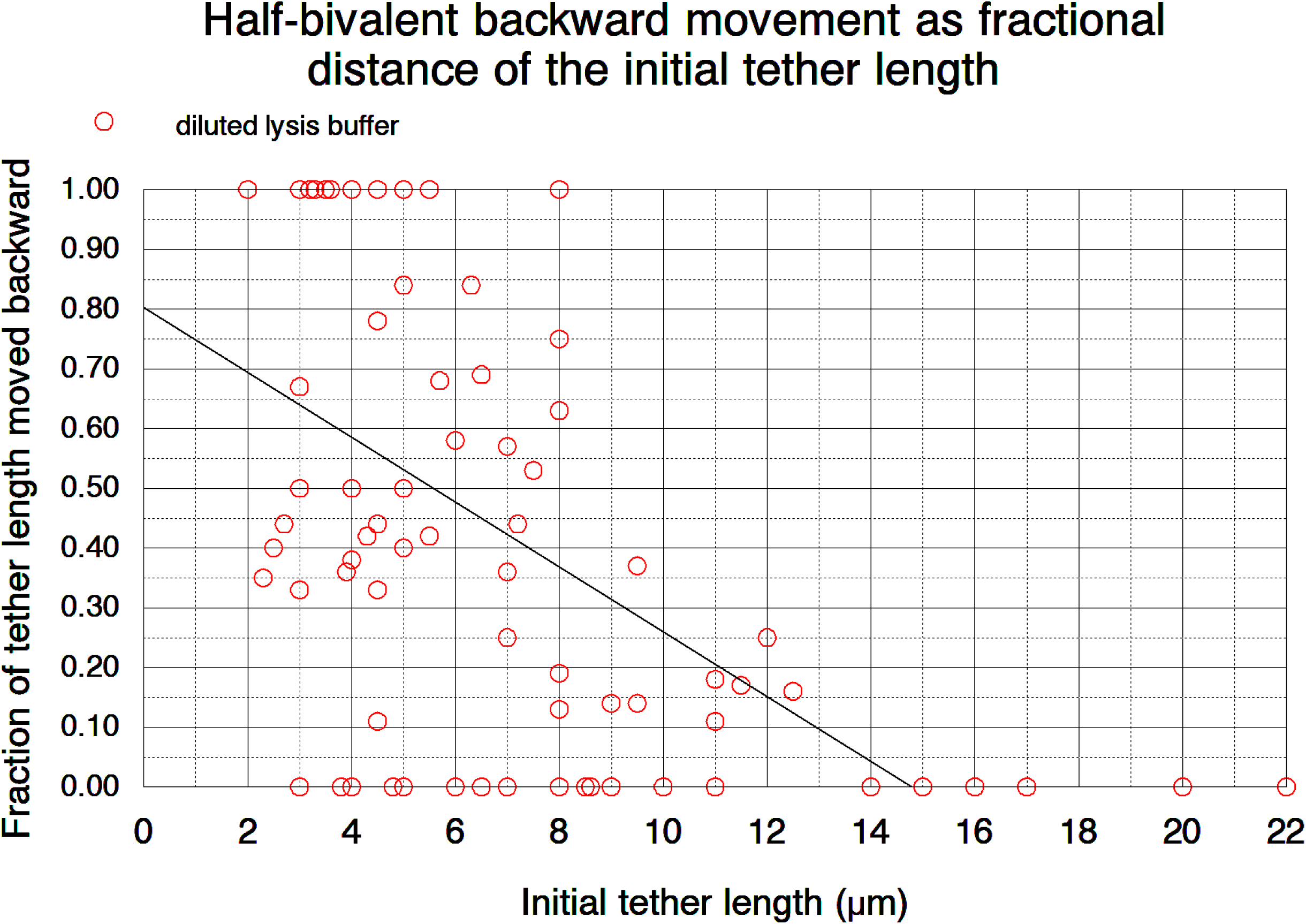
Distances moved backwards by half-bivalents as fractions of their initial tether length. Cells were treated with lysis buffer are considered in aggregate: different lysis buffer dilutions were included. 100 half-bivalent pairs were examined in total. Spearman’s rank correlation (r_s_) was calculated to assess the relationship between fractional distance moved backward and initial tether length. A negative correlation was found; with correlation coefficient (rs) of -0.610, and with alpha level ≤ 0.01, the p value was 1.62 x **10^-11^**

**Figure 8C.**
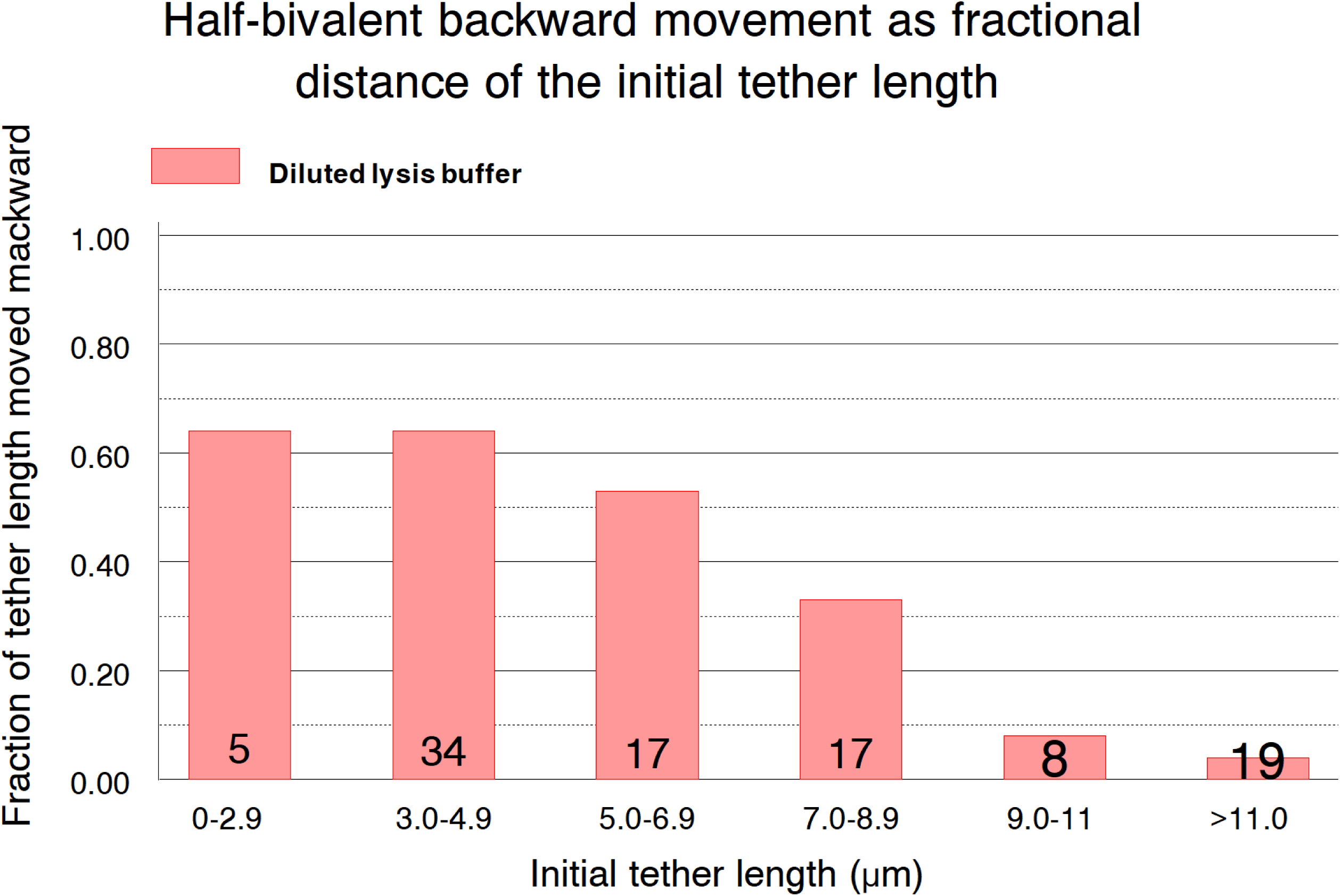
Scatter plot data from Figure 8A were grouped into categories of tether lengths and plotted against the average fractional distances moved backward. 100 half-bivalent pairs were examined in total; the numbers tested in each category are written on the respective bars.

**Figure 8D.**
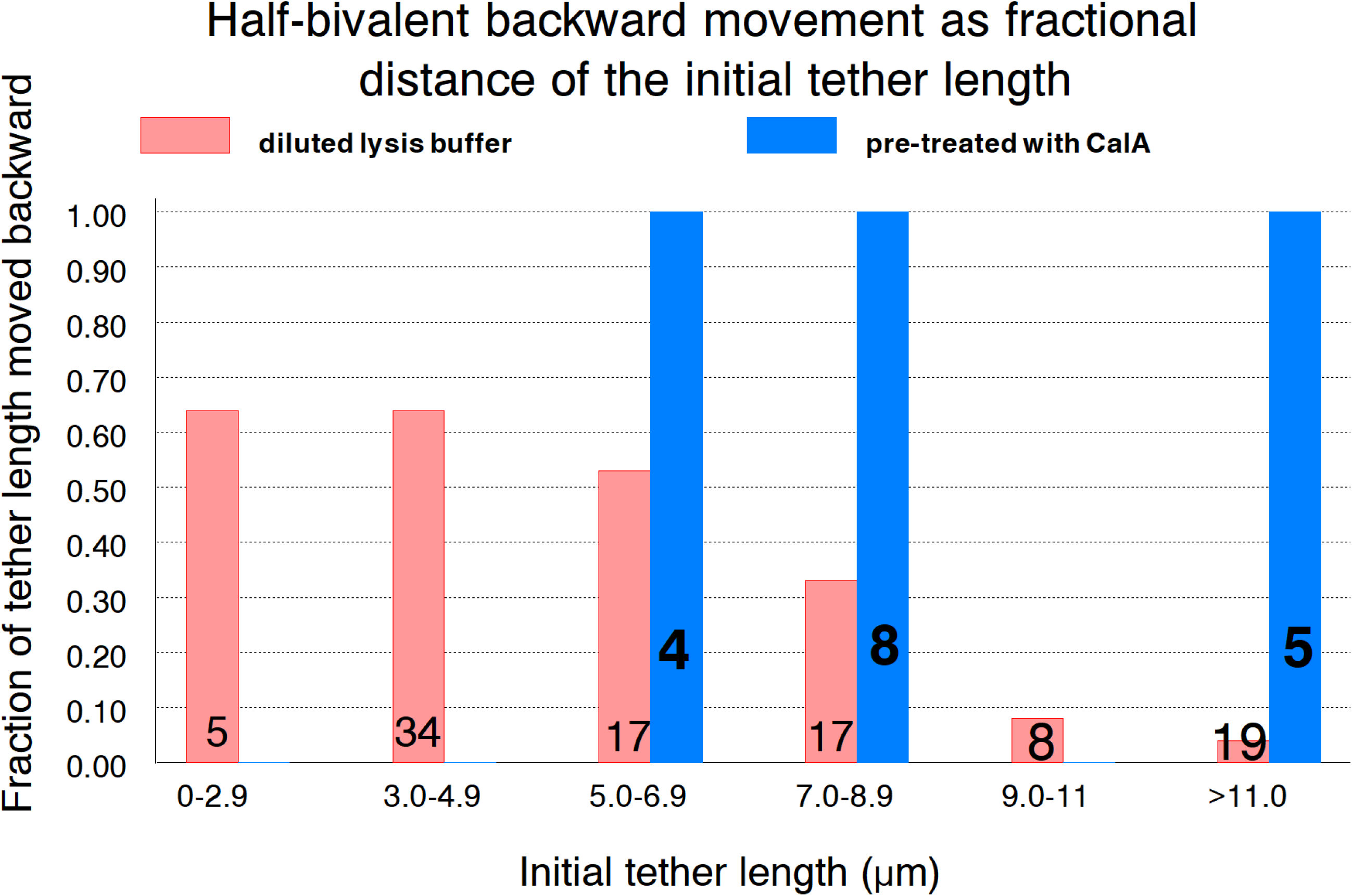
Scatter plot data from Figure 8A grouped into categories of tether lengths were plotted against the average fractional distances (of initial tether length)that the half-bivalents moved backward. Cells were treated solely with dilute lysis-buffer (pink bars) or were pre-treated with CalA before being treated with dilute lysis-buffer (blue bars). 117 homologous pairs were examined in total. The numbers in each grouped category are written on the respective bars.

**Figure 9.**
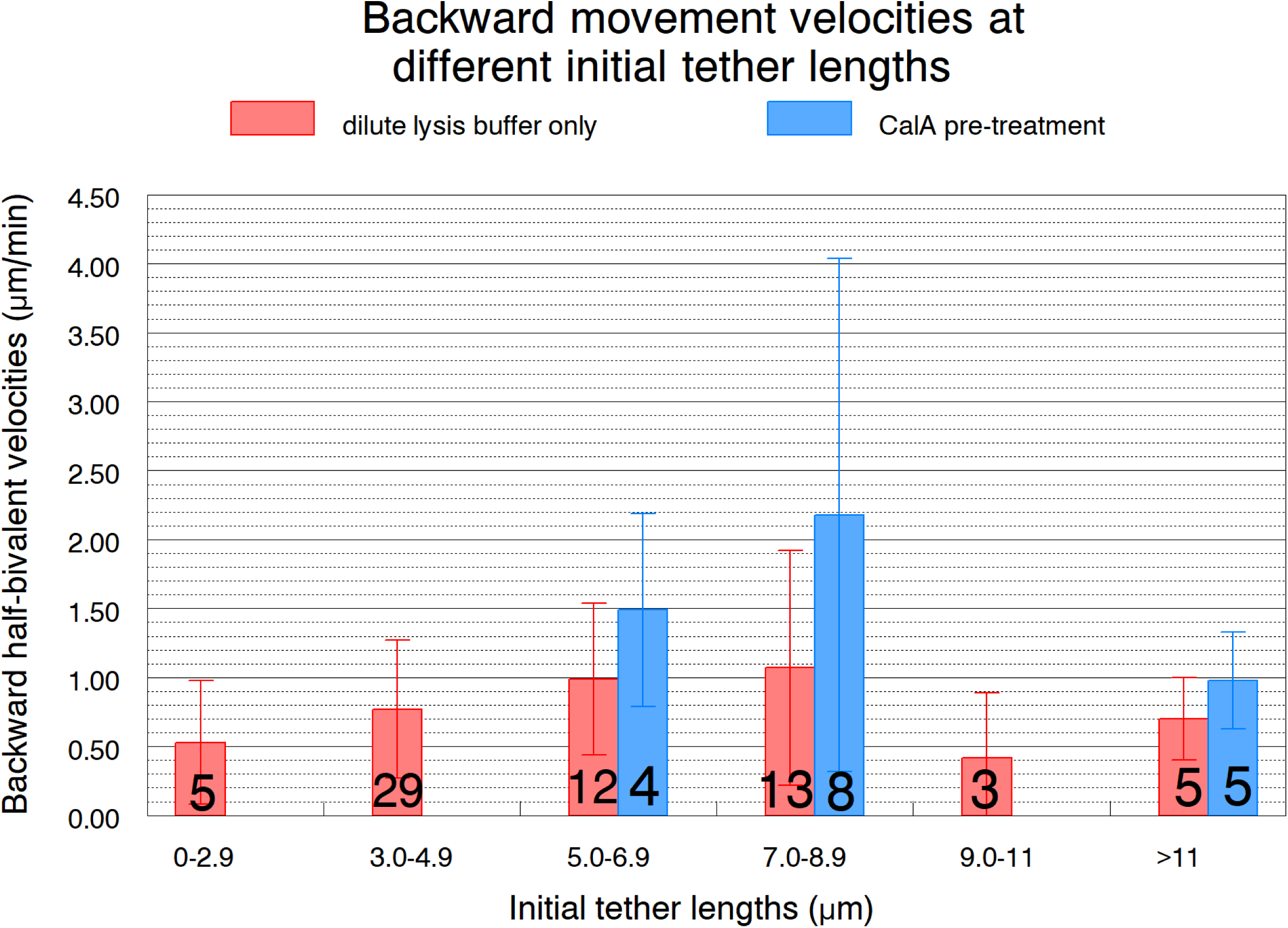
Average backward movement velocities. Average half-bivalent velocities and standard deviations (capped bars) are shown at different tether lengths during treatments with lysis-buffer only (pink bars), or after CalA pre-treatment followed by lysis-buffer (blue bars). 84 half-bivalent pairs were examined in total. The numbers in each tether length category are written on the respective bars. Student’s t-test was performed on all data sets and all differences were statistically insignificant (alpha level ≥ 0.01). Lysis buffer concentrations were used in aggregate, so any given tether length may have cells treated with different lysis buffer dilutions.

CalA applied at the beginning of anaphase (in order to maintain phosphorylated elastic tethers) followed by later treatment with dilute-lysis buffer (containing CalA), resulted in all tethers shortening 100% of their initial length no matter what the initial tether lengths were (Figure 8A, Figure 8D). That is, in cells that were pre-treated with CalA, tethers have their elasticity fully preserved at the time that lysis buffer (containing CalA) was added, so the tethers shorten the full length of the tethers when poleward movements are stopped. This further corroborates the conclusion that lysis treatment does not alter tether elasticity and functioning.

We also measured velocites of the backward moving chromosomes. Forer *et al*. (2021) reported that there were no statistically significant differences in backward velocities of chromosomal arm fragments cut at different tether lengths in both untreated and CalA-treated cells; the arm fragments moved at the same speeds in non-treated cells and Cal-A -treated cells no matter the tether lengths. The same holds true in our experiments for backward chromosome movement in cells treated with diluted lysis-buffer, as shown in Figure 9. Using Student’s t-test, there were no statistically significant differences between the velocities at the different tether lengths.This also was true for cells pre-treated with CalA, and the velocities were not statistically significantly different from cells not pre-treated with CalA. Therefore, tethers of different lengths cause backward movements with statistically similar velocities in all partially lysed cells, the same as arm-fragment movements in non-lysed cells, further corroborating that tether-induced movements in partially-lysed cells are similar to movements of arm fragments in non-treated cells.

On the other hand, movements of arm fragments are considerably faster than movements of whole chromosomes in partially lysed cells or in living cells treated with CalA, as shown in Table 3. We suggest that the chromosomes move slower than arm fragments because the chromosomes remain attached to kinetochore microtubules.

**TABLE 3.** Comparisons of arm fragment and chromosome speeds.

|  | **Chromosome**<br>***arm fragment***<br>**average speeds** | ***Half-bivalent* average**<br>**backward speeds**<br>**in partially-lysed cells** | ***Half-bivalent* average**<br>**backward speeds in**<br>**living cells** |
| --- | --- | --- | --- |
| **Not –treated cells** | 4 -7 µm/min<br>{Forer et al., 2021} | 0.4-1.1 µm/min<br>{Figure 9} | N/A |
| **Cal-A treated cells** | 6 -10 µm/min<br>{Forer et al., 2021} | 1-2.2 µm/min<br>{Figure 9} | 2.1 µm/min<br>{Table 2 of Fabian et al.<br>(2007a)} |

#### (5) After treatment with dilute lysis buffer the backward moving chromosomes remain attached to their kinetochore microtubules

We tested whether kinetochore spindle microtubules remain attached to backward-moving chromosomes in cells treated with dilute lysis buffer. We stained control and 6:1000 lysis-buffer-treated cells for tyrosinated tubulin and for acetylated tubulin (Figure 10A). Kinetochore microtubules in crane-fly spermatocytes are the only acetylated microtubules in crane-fly spermatocytes, but the acetylation does not stain near the kinetochore because that is where the tubulin monomers enter the microtubules and it takes time for newly polymerised microtubules to become acetylated (Wilson and Forer 1989, 1997; Wilson et al., 1994). Kinetochore microtubules continue to incorporate tubulin during anaphase (LaFountain et al., 2004) so the “gap” in acetylation at the kinetochores (Figure 10) persists during anaphase (Wilson et al., 1994; Wilson and Forer., 1997), though it may get smaller as the autosomes near the poles. Acetylated kinetochore microtubules are present after treatment of cell with 6:1000 dilution lysis buffer, but there seems to be no “gap” in acetylation near the kinetochore (Figure 10). We interpret this to mean that the partial lysis interferes with incorporating tubulin into the kinetochore microtubules so that the recently incorporated tubulin monomers do not move to the poles but rather sit where they were incorporated and become acetylated there. Regardless of this possible interpretation of the possible absence of the ‘gap’ in acetylation at the kinetochore, acetylated kMTs remain attached to chromosomes during lysis buffer treatment and the acetylated kinetochore microtubules may resist the backward chromosomal movements. This would lead to backward chromosome movements being held back and thus be slower than movements of unhindered chromosome arm fragments.

**Figure 10(A-C):**
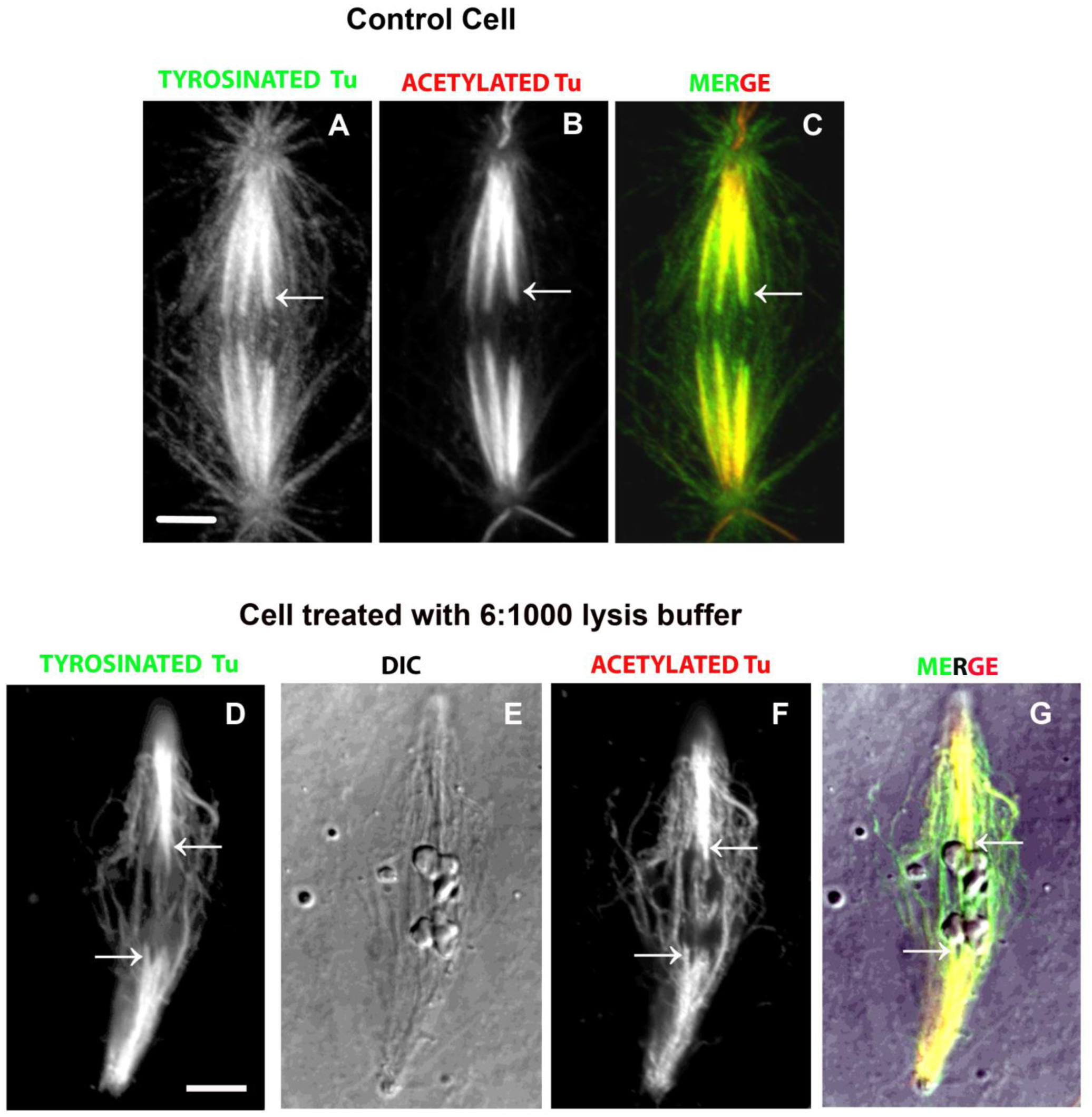
Metaphase spindle in a control cell, and **(D-G)** an anaphase spindle in a cell treated with 6:1000 dilution lysis buffer. 5µm scale bars are shown in **A** and **D**. Both cells were dual labelled with YL1/2 antibody against tyrosinated tubulin, as labelled, and 611B-1 antibody against acetylated tubulin, as labelled. The control cell images **(A-C)** are the superposition (Z-stack) of 15 confocal optical slices. The kinetochore microtubules are the only spindle microtubules that contain acetylated tubulin and the acetylated tubulin does not occur near the kinetochore itself (**A-C)**, as discussed in Wilson et al. and Forer (1989) and Wilson et al., (1994, 1997): tubulin is incorporated into kinetochore microtubules at the kinetochore and treadmills to the pole. Since there is a time-lag between incorporation and acetylation this gives rise to a gradual increase in acetylation toward the pole as illustrated in Fig. 10C, the merged image: the tubulin stain near the kinetochore indicated by the arrow (as well as by arrows in **10A** and **10B**) is tyrosinated tubulin, green, and the overlap towards the pole (yellow) indicates where the kinetochore microtubules are acetylated. Figures 10 **(D-G):** spindle staining after an anaphase cell was treated with dilute lysis buffer and the chromosomes moved backward. The images in **D-F** are the superposition (Z-stack) of 3 confocal optical slices. The DIC image illustrates the backward-moving chromosomes. The arrows in the tyrosinated tubulin **(D**) and acetylated tubulin **(F)** images point to kinetochore microtubules. The same kinetochore microtubules are seen in the merged image **(G)**, which illustrates that prominent non-kinetochore microtubules remain after treatment with dilute-lysis buffer (green images), but there is no gap in staining at the kinetochore: the yellow (overlapping stain) in the kinetochore microtubules extend up to the kinetochores.

#### (6) The lysis buffer component most likely responsible for its effects is Igepal

We tested each component of the lysis buffer to see which was/were responsible for stopping anaphase chromosome poleward movement while maintaining tether function. Each lysis component was tested separately on anaphase cells. As evident in Figure 11A, Igepal consistently stopped anaphase segregation when used at the concentration found in the 6:1000 lysis buffer dilution. Chromosomes pairs moved backward after the poleward movement stopped (Figure 11B), very similar to effects of diluted lysis buffer treatment (Figure 7). Thus, shorter tethers cause more frequent backward movements than longer tethers in Igepal-treated cells, similar to the effects of diluted lysis buffer. Further, when Ringers solution washed out Igepal, the treated cells completely recovered (in 20 out of 20 pairs of half-bivalents) within 5-7 minutes of washout, completely finishing anaphase in a normal manner. This is a higher frequency of reversing the buffer effects than the 13/46 recovery rate for washing out 6:1000 dilutions of lysis buffer (Figure 6; Table 2).

**Figure 11A.**
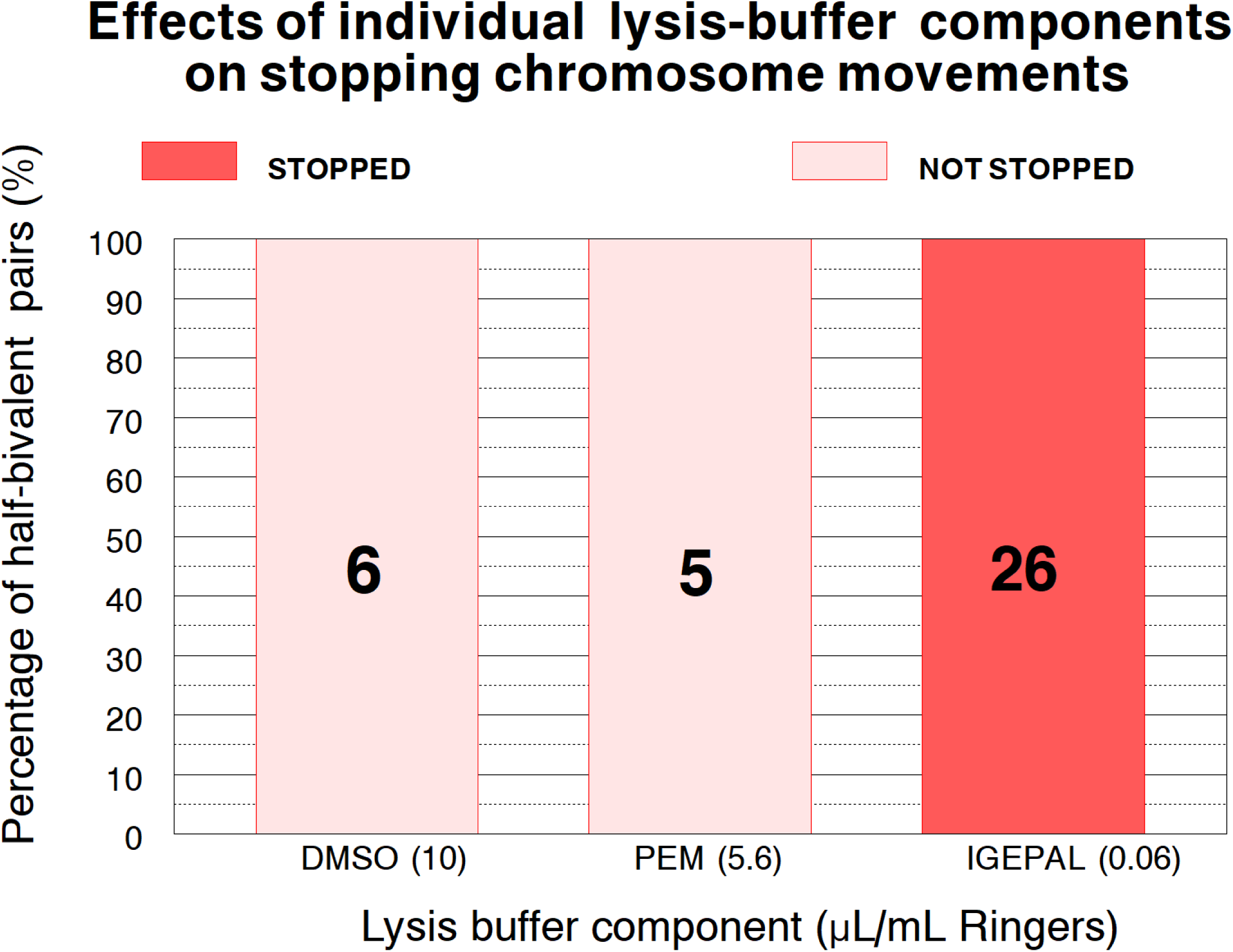
Components of lysis buffer were tested individually at their concentrations present in 6:1000 diluted lysis buffer. We measured whether the individual components stopped anaphase chromosome movement. The concentration of each component in µL/mL Ringers solution is written in brackets along the abscissa. 37 half-bivalent pairs were examined; the numbers tested with each lysis component are written on the respective bars.

**Figure 11B.**
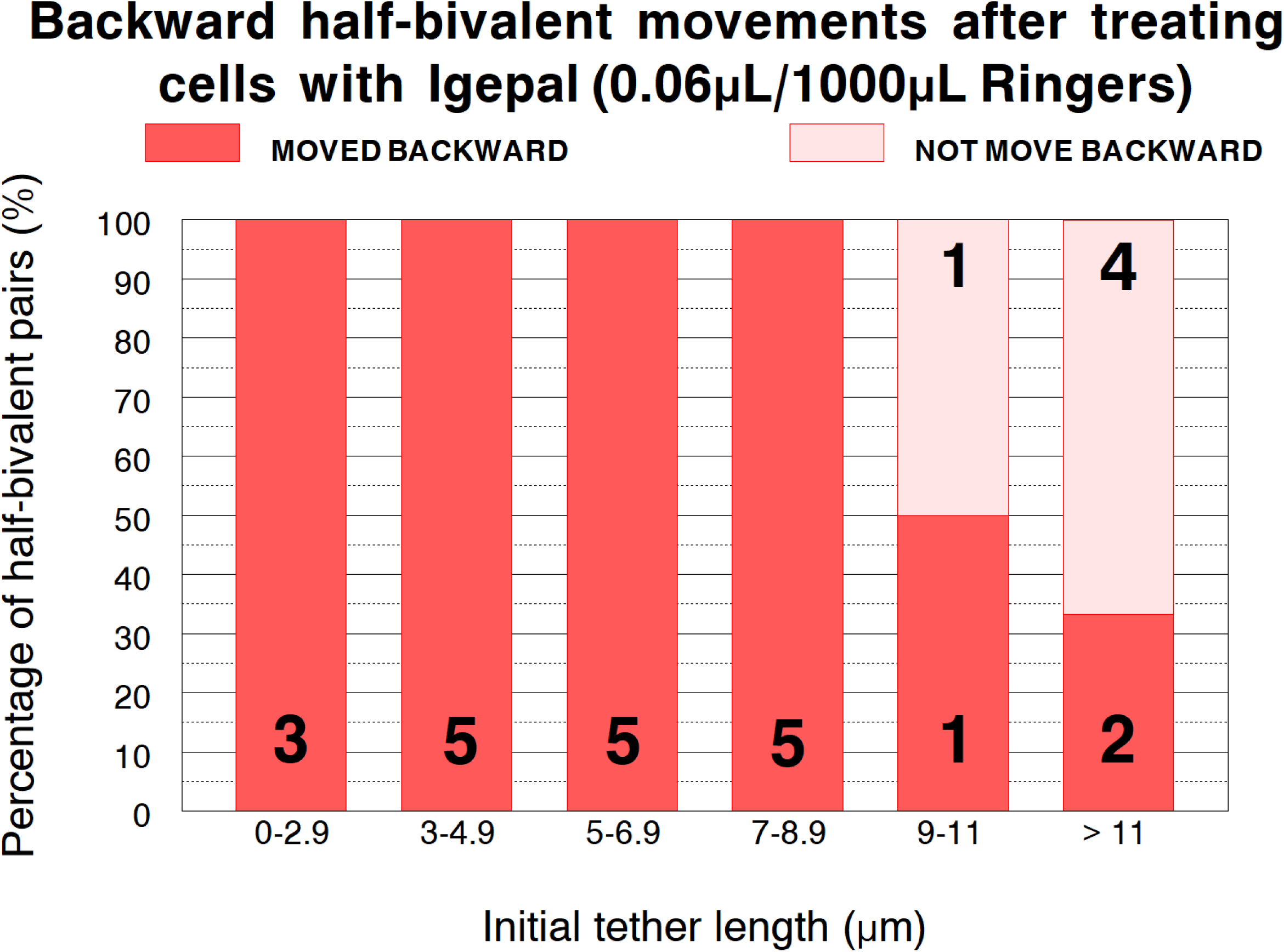
Frequency of backward movements at different tether length categories in IGEPAL treated cells. 26 half-bivalent pairs were examined. The numbers tested in each category are written on the respective bars. When comparing frequencies using Igepal with frequencies using diluted lysis buffer (Figure 7), the probability that differences at tether lengths of 3-5, 5-7, and 7-9 µm are due to chance are 0.45, 0.18, and 0.20 respectively (alpha level ≤ 0.01). Therefore, these differences all are statistically insignificant.

**In sum**, in every criterion measured, backward movements that occurred after partial lysis of cells caused by Igepal or diluted lysis buffer were the same as the backwards movements of arm fragments severed by lasers from anaphase chromosomes. Except for velocities: arm fragment moved faster than the chromosomes did. The most likely reason for that ‘discrepancy’ is that movements of chromosomes had to work against the acetylated microtubules which remained attached to the kinetochores after treatment with lysis buffer.

## DISCUSSION

The main conclusion of this research is that elastic tethers function normally after anaphase chromosome movements are stopped by dilute lysis buffer or dilute Igepal. This conclusion assumes that all the backward chromosomal movements seen during lysis-induced anaphase arrest are due to elastic tether functioning. The main evidence that supports this assertion is that the backward chromosome movements that occur after treatment with dilute Igepal or in dilute lysis buffer have characteristics similar to movements of arm fragments, and to movements of chromosomes when cells are treated with CalA, which we know are due to tethers, Forer et al., (2021). The following are the similarities. *(1)* The backward chromosomal movements are led by telomeres moving toward their partner’s telomeres while the kinetochores trail behind (Figures 3A, 4A); *(2)* Shorter tethers more consistently cause backward movements of chromosomes than longer tethers (Figure 8); *(3)* Shorter tethers cause backward movements of chromosomes over greater fractional distances than longer tethers (Figures 9A, 9B); *(4)* backward chromosome movement velocities caused by tethers of different lengths are statistically not significantly different (Figure 10); *(5)* Backward movement velocities of chromosomes in cells treated with Cal A are the same as those in cells not treated with CalA; *(6)* after pretreating cells with CalA in early anaphase the tethers remain elastic such that when lysis is added afterwards the partner half-bivalents move backward until the telomeres meet (Figures 8, 9D). All these points indicate that it is most likely elastic tethers which exert anti-poleward forces on chromosomes to cause their backward movements after anaphase chromosome movements stop due to Igepal or to dilute lysis buffer. This in turn indicates that elastic tethers function normally when the anaphase spindle stops moving chromosomes, i.e., when the spindle is non-functional, that they are two separate force systems.

The only difference between the action of tethers in partially-lysed cells and movement of chromosome arm fragments is the velocities of the movements: arm fragments move considerably faster than chromosome movements in partially-lysed (Table 3). Since the backward chromosome velocities in partially-lysed cells are similar to the velocities of backward moving chromosomes in in living cells after CalA treatment (Table 3) and since acetylated microtubules continue to extend between kinetochores and poles after cells are partially lysed (Figure 10), it seems reasonable to suggest that the differences in speed are due to the acetylated microtubules acting as an anchor, presenting forces in the direction opposite to the forces from tethers and holding back the chromosomes from freely moving backward, forces that are not experienced by cut arm fragments (Anjur-Dietrich *et al.,* 2021).

The dilute lysis-buffer and Igepal, were used at concentrations lower than those that would completely lyse the cells and release its contents; that is, they were used at concentration that partially lyse cells but that create holes in the intact cell membrane for molecules to leave and to enter the cell (Amidzadeh *et al*., 2014; Behbehani *et al*., 2014). Because normal anaphase ensues and is completed normally after Ringers solution washes out the detergent, the effects are completely reversible except at the highest concentrations of lysis buffer (6/1000) [Figures 6, Table 2]. Thus while it is apparent that Igepal or lysis buffer caused the spindle to become non-functional, no longer able to produce poleward forces, the proteins and other components involved have not been denatured or otherwise destroyed, otherwise the effects would not have been reversible. The partial lysis might block anaphase because dilute Igepal detergent punches holes in a few areas of the cell membrane and the pathways thus created could allow some intracellular proteins playing vital roles in anaphase to move out of the cell, thereby arresting anaphase (e.g., Seddon *et al*., 2004; Churchward *et al*., 2005; Sinha *et al*., 2017).

Most theories and textbooks explain anaphase chromosomal segregation as being driven primarily by microtubules and their associated motor proteins (Coue *et al*., 1991; Sharp *et al*., 2000; Rogers *et al.,* 2005). Some studies, however, suggest that actin, myosin, titin, and other spindle matrix proteins as valuable contributors to the anaphase mechanics (Forer *et al.,* 2003; Fabian *et al.,* 2007a, 2007b; Woolner et al., 2008; Johansen *et al*., 2011; Sheykhani *et al.,* 2013; Mogessie and Schuh, 2017; Forer *et al*., 2018).

These and other studies suggest that the anaphase spindle apparatus is a cytoskeletal structure composed of a variety of protein molecules interacting with each other and with the chromosomes via complex mechanisms to achieve successful chromosomal segregation (Silverman-Gavrila and Forer, 2000; Fabian and Forer, 2005; Pimm and Henty-Ridilla, 2020). Regardless of which story of force production turns out to be correct, data presented here show that we need in addition to account for forces produced by tethers on anaphase chromosomes. It might be worth pointing out that our experiments were carried out on cells in which anaphase occurs solely without spindle elongation: in crane-fly spermatocytes autosome anaphase movements are due solely to shortening of kinetochore fibres, with the poles remaining at fixed positions. We expect that when cells are partially lysed tethers will still be elastic but we do not know how the elasticity will be manifest in the possible presence of pole elongation (Anaphase B).

To summarize, our experiments show that partially lysing cells arrests anaphase chromosome movements but does not affect the function of the tethers that connect the separating anaphase telomeres of each chromosome pair during anaphase. The elastic tethers produce backward forces on chromosomes in partially lysed cells with the same characteristics as they do in non-lysed (control) cells. Therefore, partially lysed cells might be able to be further developed into an *in vitro* system for studying tethers and other spindle components to better understand their contribution to normal cell division.

## ACKNOWLEDGEMENTS

The work reported herein was supported by grants to AF from the Natural Sciences and Engineering Research Council (NSERC) of Canada to AF and an NSERC student Fellowship to AA. Some of the material in this article was presented by AA to York University as part of her thesis for an M.Sc. degree,

